# Robust capability of renal tubule fatty acid uptake from apical and basolateral membranes in physiology and disease

**DOI:** 10.1101/2022.07.04.498762

**Authors:** Ryo Kawakami, Hirofumi Hanaoka, Ayaka Kanai, Hideru Obinata, Daisuke Nakano, Hidekazu Ikeuchi, Miki Matsui, Toshiyuki Matsuzaki, Rina Tanaka, Hiroaki Sunaga, Sawako Goto, Hiroki Matsui, Norimichi Koitabashi, Keiko Saegusa, Tomoyuki Yokoyama, Keiju Hiromura, Akira Nishiyama, Akihiko Saito, Motoko Yanagita, Hideki Ishii, Masahiko Kurabayashi, Tatsuya Iso

**Author notes:** Corresponding author (TI).

## Abstract

Excess lipid accumulation is associated with obesity-related chronic kidney disease, but the mechanisms of fatty acid (FA) uptake have been poorly understood. To this end, we investigated how FAs are taken up by tubular epithelial cells (TECs) in mice by using in vivo FA tracing and histological methods. Immunohistochemistry showed that CD36, which is a well-known FA transporter, was abundantly expressed on the basolateral side of proximal TECs (PTECs). The uptake of ^125^I-BMIPP (a radiolabeled FA tracer) was significantly reduced in CD36-knockout kidneys at 1 min after injection. In vivo imaging with multiphoton microscopy revealed that BODIPY-C_12_ (a fluorescence-labeled FA tracer) accumulated on both the basolateral and apical sides of PTECs. Numerous lipid droplets accumulated in PTECs after accelerated lipolysis. Furthermore, PTEC-specific injury via diphtheria toxin (DT) injection in transgenic mice expressing the DT receptor resulted in a compensatory increase in lipid accumulation in downstream TECs. Importantly, urinary FAs were undetectable, even in mice and humans with remarkable albuminuria. Our data demonstrate that renal TECs take up FAs from blood (CD36-dependent) and primary urine (CD36-independent) and can store excess FAs as neutral lipids. The results further show that renal tubules have hitherto largely unappreciated mechanisms by which the excretion of FAs into the urine is avoided.

## Introduction

Prevalent chronic kidney disease (CKD) parallels the epidemics of obesity and diabetes. Lipid accumulation in the glomeruli and tubular epithelial cells (TECs), as well as the resulting lipotoxicity, have been suggested to be associated with glomerulosclerosis and tubulointerstitial pathologies in animal models and human kidney diseases (Bobulescu, 2010, Wahl et al., 2016). The kidney is one of the most energy-demanding organs. The resting energy expenditure of the kidney is the highest among organs, comparable to that of the heart (440 kcal/kg/day) (Gallagher et al., 1998). Two-thirds of the oxygen consumption for ATP synthesis is accounted for by fatty acid (FA) oxidation (Mount et al., 2015). FAs are supplied to the kidney as either albumin-bound FAs or as FAs released from the triacylglycerides (TGs) contained in TG-rich lipoproteins, such as chylomicrons and very-low-density lipoproteins (Iso and Kurabayashi, 2017). The kidney requires a large amount of energy to reabsorb nutrients and small proteins in primary urine, as well as to regulate the balance of electrolytes, fluid volume, and acid-base homeostasis (Bhargava and Schnellmann, 2017). In particular, a considerable amount of ATP is consumed by Na^+^-K^+^ ATPases to generate ion gradients across the basolateral membrane in the proximal tubules (PTs), the thick ascending limbs of Henle’s loop (TALs), and the distal convoluted tubules (DCTs) (Soltoff, 1986, Tian and Liang, 2021). Consistently, epithelial cells in these segments are rich in mitochondria for FA utilization (Bhargava and Schnellmann, 2017). However, despite the high demand for FAs, little is known about the route and molecular mechanisms underlying FA uptake in the kidney.

In contrast to the mechanisms underlying FA uptake in the kidney, those mechanisms in the heart, muscle, and adipose tissues are well established. FA uptake in tissues is regulated by several crucial molecules, including lipoprotein lipase (LPL) and CD36 (Pi et al., 2018, Abumrad et al., 2021, Hasan and Fischer, 2021, Iso and Kurabayashi, 2021). At the luminal surface of capillary endothelial cells, LPL hydrolyzes the TGs contained in TG-rich lipoproteins in plasma, thus resulting in the liberation of FAs. Liberated FAs or albumin-bound FAs from adipose tissues are taken up by parenchymal cells via CD36, which is a single-chain membrane protein. Although lipid accumulation occurs in both the heart and the kidney during prolonged fasting, the mechanisms for each organ are distinct (Scerbo et al., 2017, Trent et al., 2014). Cardiac lipid droplets are dependent on LPL activity (Trent et al., 2014), whereas kidney TG accumulation is highly associated with serum FA levels (but not LPL activity) (Scerbo et al., 2017). Importantly, in mice lacking LPL or CD36, lipid deposits are increased in the kidney but not in the heart, even during fasting (Scerbo et al., 2017, Trent et al., 2014). These findings strongly suggest that the kidney has unique and unrevealed mechanisms/molecules for taking up FAs and for depositing TG droplets.

The majority of FAs in the plasma are carried by albumin. Recent studies have demonstrated that a certain amount of plasma albumin is passively filtered through the glomerulus, although the calculated filtration rate is very wide (ranging from 0.5–3.5 to 200 g/day) (Bobulescu, 2010, Dickson et al., 2014, Comper et al., 2008, Birn and Christensen, 2006). It has been reported that filtered albumin is reabsorbed by several candidate molecules, such as megalin and cubilin, which is a multiligand receptor complex located at the brush border of the apical membrane (Dickson et al., 2014). Together with albumin reabsorption via the megalin/cubilin complex, albumin-bound FAs may also be reabsorbed from the apical membrane. Thus, both albumin and albumin-bound FAs are suggested to be filtered through the glomerulus and reabsorbed from the apical membrane of proximal TECs (PTECs). However, little attention has been given to how FA uptake from the apical and basolateral sides is coordinated under physiological and pathological conditions. In this study, we addressed the issue of how FAs are taken up by tubular epithelial cells in mice lacking CD36 or in mice whose PTECs are conditionally injured upon diphtheria toxin injection. We also examined FA excretion in urine in PTEC-injured mice and CKD patients with massive proteinuria. Based on our results, we demonstrated three essential mechanisms. First, FAs are taken up from both the basolateral side via CD36 and the apical side independently of CD36. Second, lipid accumulation occurs along the structure of the nephron when serum FA levels are increased. Third, FAs in primary urine are completely reabsorbed while passing through the nephron, thus resulting in no detectable FAs in urine. Thus, renal tubules have elaborate mechanisms for taking up FAs and for avoiding FA excretion into the urine.

## Results

### Neutral lipids predominantly accumulate in the cortex of the kidney after prolonged fasting

Renal lipid accumulation is closely associated with an increase in serum FA levels (Scerbo et al., 2017). We first examined the relationship between serum FA levels and TG content in the kidney after fasting. Serum FA levels were elevated 16 and 24 h after fasting, accompanied by an increase in TG content in the kidney (Fig 1A). TG content in the kidney was positively associated with serum FA levels (Fig 1B), as reported (Scerbo et al., 2017). Oil red O staining revealed that neutral lipids predominantly accumulated in the cortex 16 h after fasting (Fig 1C). Thus, fasting-induced lipolysis results in TG accumulation in the kidney.

**Figure 1.**
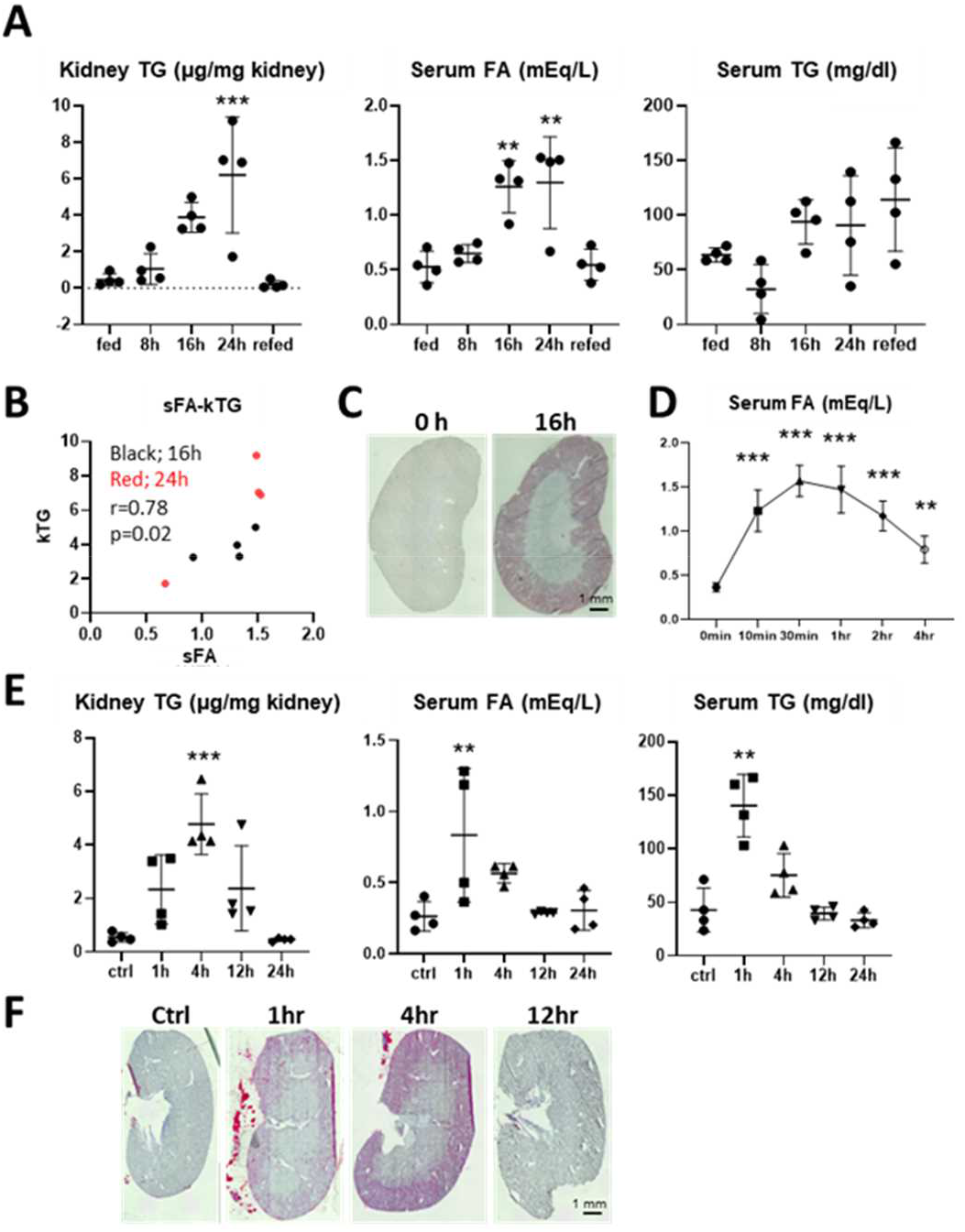
Neutral lipids accumulate in the kidney after accelerated lipolysis. (A-C) Wild-type mice were either given food ad libitum (fed) or fasted for 8, 16, or 24 h. Refed indicates a 24-h fast followed by a 24-h feed. (A) Lipids were extracted from the kidneys for triglyceride (TG) measurement. Fatty acids (FAs) and TGs in serum were measured. (n = 4) *p < 0.05, **p < 0.01, ***p < 0.001 vs. fed group. (B) There is a simple correlation between serum FA levels and TG content in the kidneys from 16 h- and 24 h-fast groups. Pearson’s coefficient (r) and the p value are shown. (C) Representative images of oil-red O staining for detecting neutral lipid accumulation in the kidney. Scale bar, 1 nun. (D-F) β3 adrenergic agonists. CL316.243, were intraperitoneally injected into wildtype mice. (D) Time course of serum FA levels after intraperitoneal injection (ip) of CL316.243. (n = 5) **p < 0.01. ***p < 0.001 vs. 0 min control (Ctrl). (E) Time course of TG content in the kidney, serum levels of FAs and TGs after CL316.243 ip. (n = 4) **p < 0.01. ***p < 0.001 vs. Ctrl. (F) Representative images of oil-red O staining of the kidneys from (E). Scale bar, 1 mm.

### Neutral lipids accumulate in the kidney by enhanced lipolysis

Adrenergic stimuli promote lipolysis, which leads to elevation of serum FA levels. After intraperitoneal administration (ip) of CL316,243, a β3 adrenergic receptor agonist, serum FA levels peaked at 30 min (Fig 1D). Neutral lipid accumulation peaked at 4 h and decreased after that (Fig 1E). When lipid deposits were maximum, most were observed in the cortex and a lesser amount in the outer medulla (Fig 1F). Before and after the peak of lipid accumulation, lipid deposits were restricted to the cortex (Fig 1F, 1 h and 12 h). Importantly, lipid accumulation was transient and did not last for a long time (Figs 1E and 1F). These findings suggest that an excess amount of FAs is esterified as TGs when FA uptake exceeds FA combustion and that accumulated TGs are diminished after the FA supply becomes low.

### Identification of lipid-accumulating cells

Next, we attempted to identify lipid-accumulating cells by simultaneous executing ORO staining and IF by using cryosections from mice 4 h after CL316,243 ip (Fig 2A). Many lipid droplets were observed in LTL-positive and AQP4-negative PTECs (S1 and S2). Fewer lipids were detected in LTL and AQP4-positive PTECs (S3), THP-positive TAL cells, and NCC-positive DCT cells. A pinkish color was marginally observed in AQP2-positive collecting duct (CD) cells (Fig 2A). When lipid accumulation was modest, accumulation was observed only in S1/S2 (Fig S1). The spatiotemporal distribution of lipid accumulation is summarized in Figure 2B. Thus, neutral lipid accumulation in S1/S2 precedes and exceeds that in the more distal nephron segments (S3/TAL/DCT). These findings raise an intriguing hypothesis that FAs for TG formation are mainly supplied from primary urine while passing through the nephron structure.

**Figure 2.**
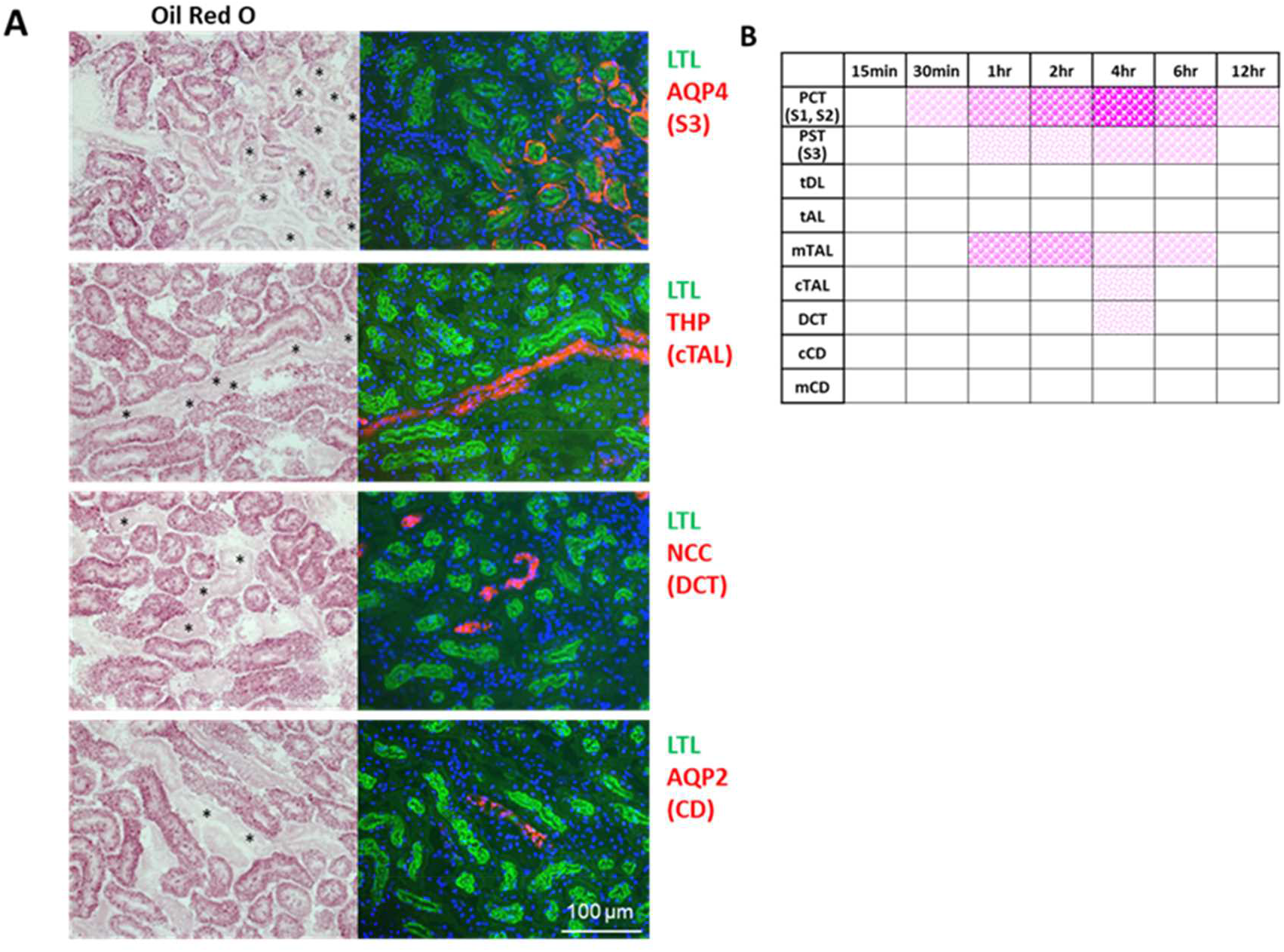
Temporal and spatial distribution of lipid accumulation after CL316,243 injections. (A) Oil-red O staining and immunofluorescence (IF) were executed simultaneously by using the kidneys from Figures ID and IE. Representative images are shown. Lotus tetragonolobus lectin (LTL), a marker for the brush border of proximal tubule epithelial cells (PTECs. green): aquaporin 4 (AQP4) for proximal straight tubules (PSTs, red); Tamm-Horsfall protein (THP) for thick ascending limbs of Henle (TALs, red); Na^+^-Cl^-^ cotransporter (NCC. red) for distal convoluted tubules (DCTs, red); AQP2 for collecting ducts (CDs, red); 4’,6-diamidino-2-phenylindole (DAPI) for nuclei (blue). The asterisks indicate red IF-positive tubules. Scale bars, 100 μm. (B) Summary of lipid distribution. The pink dots with different intensities and sizes indicate the amount and relative size of lipid droplets. PCT: proximal convoluted tubule, tDL: thin descending limb of Henle, tAL: thin ascending limb of Henle, mTAL: medullary TAL, cTAL: cortical TAL, eCD: cortical CD, mCD: medullary CD.

### FAs are taken up from both circulating blood and primary urine: the visualization of FA uptake *in vivo* by multiphoton microscopy

To explore how FAs are taken up by the kidney, *in vivo* live imaging by multiphoton microscopy was carried out with fluorescent BODIPY dodecanoic acid (BODIPY-C_12_). One minute after intravenous injection, BODIPY-C_12_ was detected in the peritubular capillary and basolateral membrane of PTECs (Fig 3A). The intensity of the BODIPY-C_12_ signal gradually increased throughout the cytosol in the late segments but not the early segments of PTs and peaked 10 min after injection (Figs 3A and S2A). In early segments of PTs, the relative intensity of BODIPY-C_12_ was higher in the brush border than in the cytosol 10 min after injection, suggesting more FA uptake from the apical side (Figs 3A and S2A). The fluorescence intensity was always higher in the late segments than in the early segments of PTs (Figs 3A and 3B). These findings suggest that FAs are predominantly taken up from circulating blood by the late segments of PTs during the early phase after injection, while they are supplied from primary urine in the early segments during the late phase. The time lag of FA uptake between the basolateral and apical membranes could be attributed to the limited glomerular sieving of FAs conjugated with albumin.

**Figure 3.**
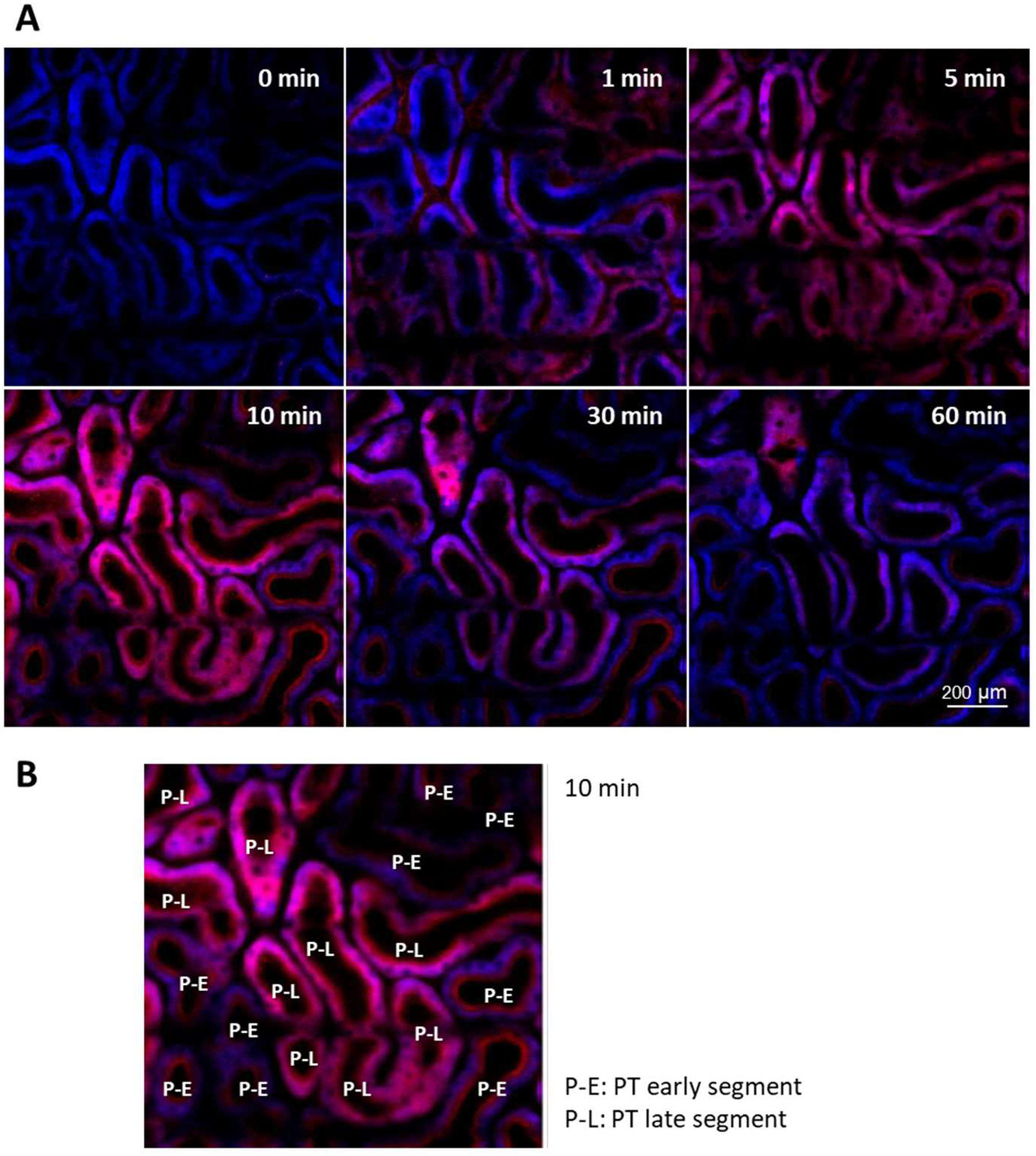
BODIPY-C_12_ accumulates on both the basolateral and apical sides of PTECs. (A, B) *In vivo* imaging of the medium-chain FA tracer 4,4-Difluoro-5-(2-Thienyl)-4-Bora-3a,4a-Diaza-s-Indacene-3-Dodecanoic Acid (BODPY-C_12_). BODIPY-C_12_ was administered to wild-type mice from the right cervical vein and observed using multiphoton microscopy. Blue and magenta represent cytosolic autofluorescence and BOD1PY-C_12_, respectively. (A) Representative images 0, 1, 5, 10, 30, and 60 min after BODIPY-C_12_ administration. Scale bars. 200 μm. (B) The early or late segments of proximal tubules (P-E or P-L. respectively) are indicated on the image at 10 min. These segments were identified by a bolus injection of FITC-inulin.

### CD36 plays an important role in FA uptake from blood in the kidney

To study the involvement of CD36 in FA uptake in the kidney, we first explored the precise distribution of CD36 expression. Interestingly, CD36 was abundantly expressed in the basolateral membranes of late segments of PTs but not in the early segments (Figs 4A, 4B, S3A, and S3B), suggesting that CD36 could play a role in FA uptake from the peritubular capillary in the late segments of PTs. The specificity of CD36 was confirmed using CD36KO kidneys (data not shown). We next assessed the uptake capacity of FAs and glucose by two lipid-consuming organs, the heart and the kidney, in CD36^-/-^ (CD36KO) mice by using ^125^I-BMIPP and ^18^F-FDG in the fasted and refed states. As reported previously, ^125^I-BMIPP uptake was markedly reduced with a compensatory increase in ^18^F-FDG uptake in CD36KO hearts in the fasted state 2 h after the injection (Fig S4A). However, ^125^I-BMIPP uptake was even higher in CD36KO kidneys than in WT kidneys, with no significant change in ^18^F-FDG uptake (Fig S4A). These findings suggest that CD36-dependent FA uptake from the basolateral membrane might be compensated by CD36-independent FA uptake from the apical membrane after glomerular filtration. We next estimated FA uptake at the earlier phase to minimize the influence of FA uptake from the apical side. Although blood levels of ^125^I-BMIPP at 1 min were much higher in CD36KO mice, the uptake at 1 min was comparable between WT and CD36KO kidneys (Fig S4B). We further perfused the mice with phosphate-buffered saline to minimize the remaining blood in the kidneys and found that ^125^I-BMIPP uptake at 1 min was significantly lower in CD36KO kidneys than in WT kidneys (Fig 4C). A reduction in ^125^I-BMIPP uptake at 1 min in CD36KO kidneys was also confirmed by autoradiography, while the uptake at 30 min was comparable (Fig 4D). We next assessed the effects of fasting and CL316,243 ip on TG accumulation. TG content in the kidney was similarly elevated in both WT and CD36KO mice 16 h after fasting, although serum FA levels were significantly higher in CD36KO mice than in WT mice (Fig 4E). It is plausible that a reduction in NFFA supply from blood due to CD36 deficiency is compensated by increased supply from primary urine due to higher levels of serum FAs. Interestingly, an increase in TG content in the kidney by CL316,243 ip in WT mice was diminished in CD36KO mice despite a similar elevation of serum FAs (Figs 4F and S4C). The reduced TG content in CD36KO kidneys treated with CL316,243 could reflect a reduction in FA uptake from the blood. Thus, FA uptake from blood is reduced in CD36KO kidneys, and the reduction is masked by compensatory FA uptake from primary urine in most cases. Collectively, our data strongly suggest that CD36 plays a role in FA uptake from blood in the kidney.

**Figure 4.**
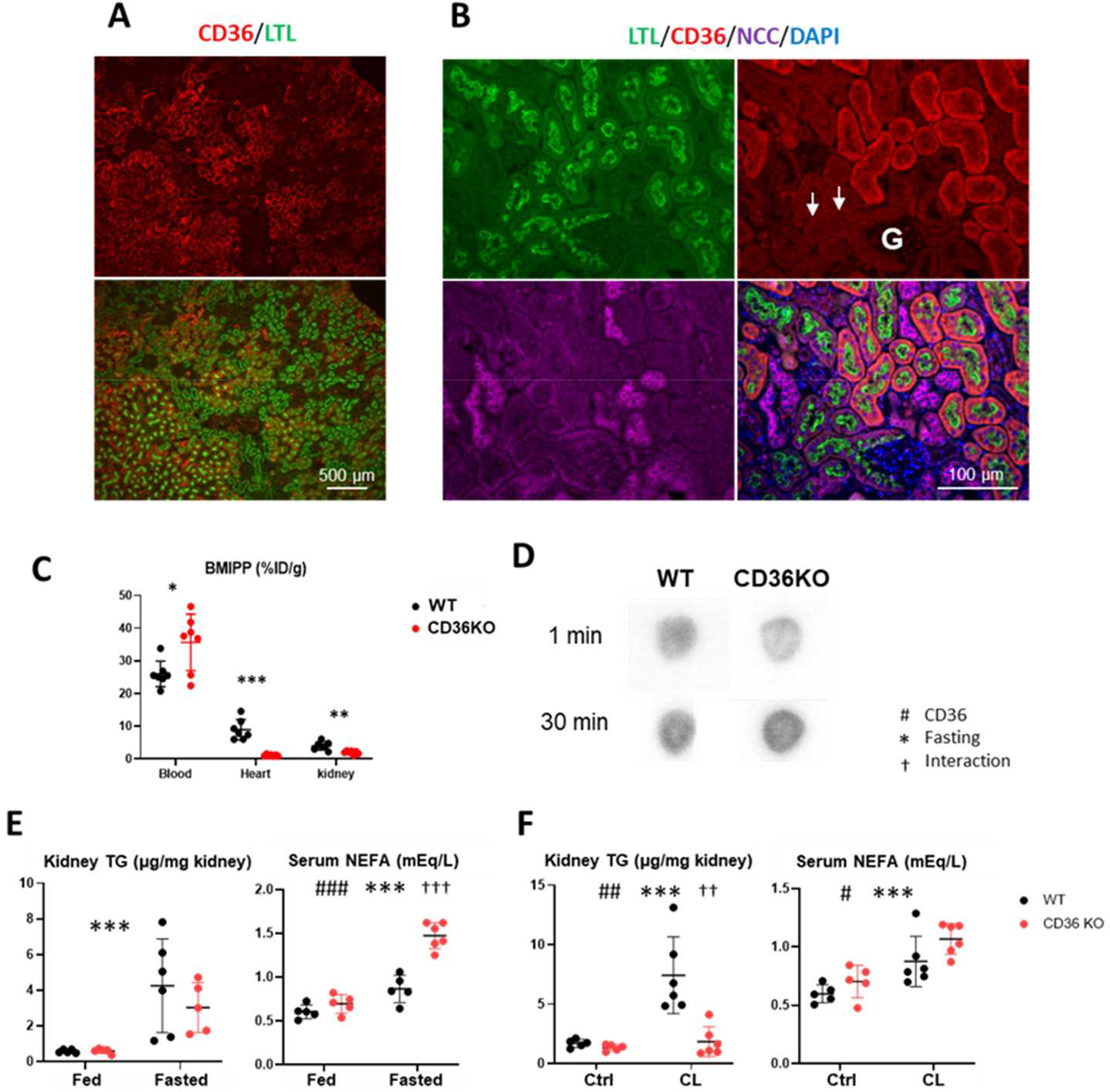
CD36 expressed in distal segments of PTECs is involved in FA uptake from circulation. (A and B) Representative images of IF. CD36 (red), LTL (green). NCC (purple), and DAPI (blue). CD36 was partially colocalized with LTL (A and B) but not with NCC (B). Note that the most proximal segments of PTECs (white arrows. SI) lacked CD36 expression. Scale bars. 100 pm. G. glomerulus. (C. D) FA uptake was assessed with 15-(p-iodophenyl)-3-(R, S)-methyl pentadecanoic acid (^125^I-BMIPP). CD36 knockout (CD36KO) and littermate control (wild-type. WT) mice fed ad libitum were used. (C) One minute after 125I-BMIPP injections, blood was drawn from the retro-orbital plexus. After systemic perfusion with phosphate-buffered saline, hearts and kidneys were isolated, (n = 7) (D) Representative autoradiographic images of short axis-sliced kidneys from (C). (E) WT and CD36KO mice were fed ad libitum or overnight fasted. Kidney TG content and serum FA levels were measured, (n = 5-6) (F) WT and CD36KO mice were fed ad libitum. Kidney TG content and serum levels of FAs were measured 4 h after CL316.243 ip. (n = 5-6) *^,#,†^p < 0.05, **^,##,† †^p < 0.01, ***^,###,† †^ p < 0.001.

### FAs in primary urine are reabsorbed by renal tubules independently of megalin

We next studied whether the lack of megalin in PTECs affects lipid reabsorption in the kidney. As reported previously (Weyer et al., 2011), megalin KO mice exhibited marked albuminuria (Fig 5A). However, FAs were undetectable in urine (Fig 5A), suggesting the complete reabsorption of FAs from primary urine by a megalin-independent process. We further explored the TG content and distribution pattern in the kidney in megalin KO mice 4 h after CL316,243 ip, which was comparable to those in WT mice (Figs 5B and 5C). Thus, it is suggested that FAs in primary urine are reabsorbed by renal tubules independent of albumin reabsorption mediated by megalin.

**Figure 5.**
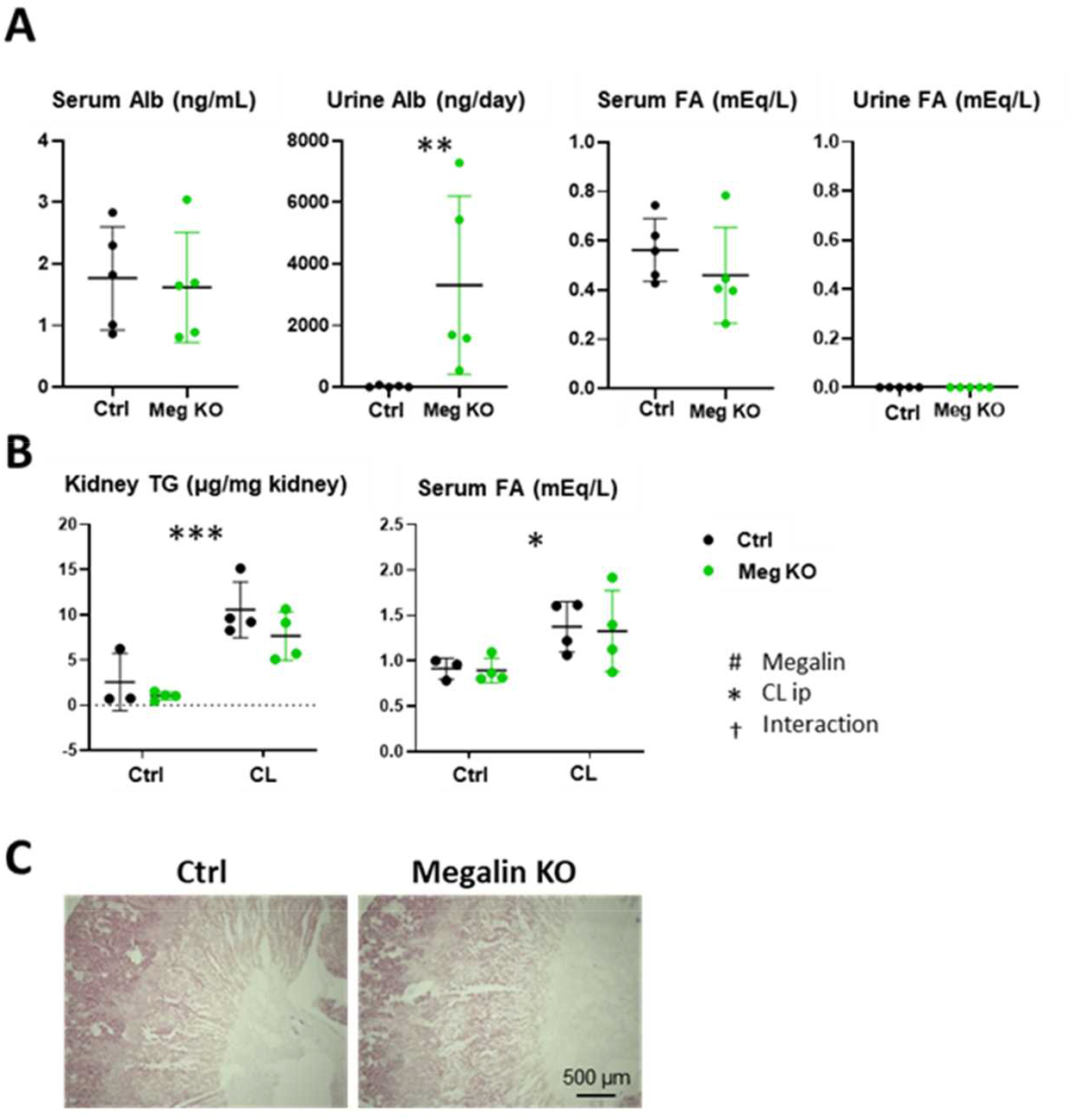
Megalin is not involved in FA reabsorption. (A) Albumin and FA levels in serum and urine were measured in megalin-knockout (Meg KO) mice and their littennate controls (Ctrl). (n = 5) ^**^p < 0.01 vs. Ctrl. (B) Kidney TG content and serum FA levels were measured 4 h after CL316.243 ip in Ctrl and Meg KO mice. (n = 4) ^*,#,†^ p< 0.05.^**,##, †^ ^†^ p < 0.01. ^***,###, †††^ p < 0 001 (C) Representative images of oil-red O staining 4 h after CL316.243 ip. Scale bars, 500 μm.

### PT injury causes lipid accumulation in the downstream nephron segments

As shown above, S1/S2 of PT are the major sites of FA uptake and TG storage. We next questioned whether PT injury affects FA uptake and TG accumulation. To address this issue, we employed PT-specific injury (PTi) mice by a system with diphtheria toxin (DT) injection and PT-specific overexpression of the DT receptor. The injury is more severe in S1/S2 than in S3 due to the difference in DT-R expression levels (Takaori et al., 2016). Marginal lipid accumulation was observed in the cortex in PTi mice without treatment (Fig 6A), suggesting a reduction in FA combustion relative to FA uptake. Four hours after CL316,243 ip, lipid accumulation in S3 was enhanced in addition to that in S1/S2 in PTi mice (Figs 6A and 6B). Neutral lipids also accumulated in subsequent segments, such as the TAL, DCT, and cortical CD (Figs 6C and S5). TG content was increased in PTi mice compared to control mice without CL316,243 ip (Fig 6D), which is consistent with marginal lipid accumulation in the cortex (Fig 6A). TG content in the kidney was comparable between control and PTi mice after CL316,243 ip (Fig 6D). These findings suggest that FA uptake and esterification were disturbed in PTECs in PTi mice and that FAs not taken up by S1/S2 were reabsorbed and stored by the subsequent nephron segments. We further found that PTi mice exhibited remarkable proteinuria, probably due to glomerular and PT injuries (Fig 6E). However, despite remarkable albuminuria, FAs were undetectable in PTi mice (Fig 6E). Taken together, these results show that it is very likely that S3/TAL/DCT/cortical CD also has a large capacity to completely reabsorb FAs in primary urine and store them as TG droplets independent of albumin reabsorption when PTs are injured.

**Figure 6.**
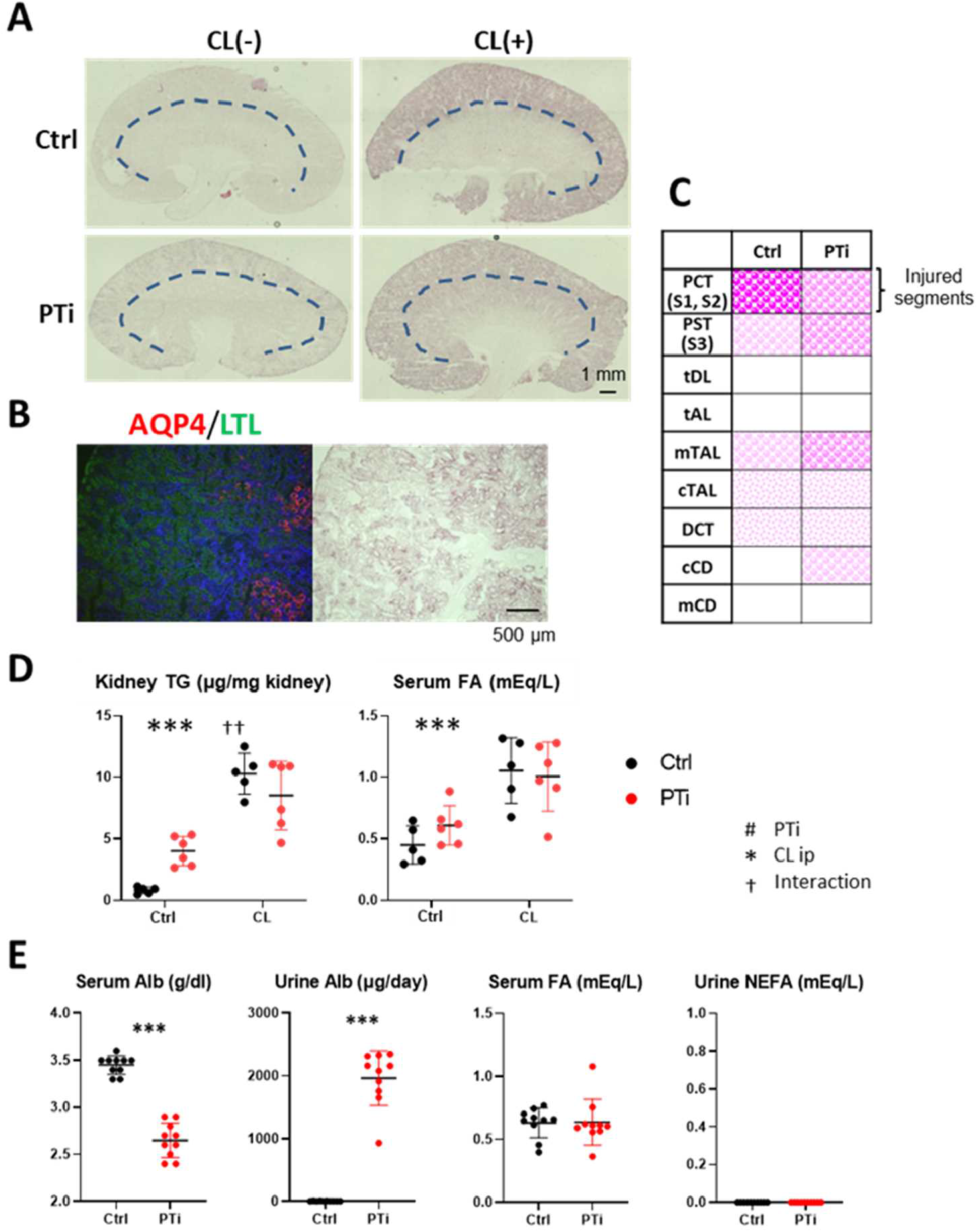
Lipid accumulation is increased in the distal nephron in proximal tubule injury mice. (A-C) CL316.243 was intraperitoneally injected into proximal tubule-specific injury (PTi) mice and their littermate controls (Ctrl). (A) Representative images of oil-red O staining 4 h after CL316,243 ip. The dashed lines indicate borderlines between the renal cortex and medulla. Scale bars. 1 mm. (B) Simultaneous staining of oil-red O and IF. AQP4. a marker for PST (red); LTL for PTECs (green); DAPI for nuclei (blue). Scale bars, 500 μm. (C) Summary of lipid distribution in Ctrl and PTi mice 4hr after CL316.243 injections. The pink dots with different intensities and sizes indicate the amount and relative size of lipid droplets. (D) Kidney TG content and serum levels of FAs were measured 4 h after CL316,243 ip in Ctrl and PTi mice, (n = 5-6) ^*,#,†^< 0.05. ^**,##,††^p < 0.01. ^* **,###,†††^p < 0.001. (E) PTi and Ctrl mice were fed ad libitum in metabolic cages for 24 h. and blood and urine were collected. Levels of albumin and FAs in serum and urine were measured. (n = 10) ^* * *^p < 0.001 vs. Ctrl.

### Urinary FAs are undetectable in patients with massive proteinuria and/or deteriorating kidney function

We next measured FAs in urine in patients with massive proteinuria and/or CKD G4–G5 (Figs 7 and S6). FAs and long-chain FAs, such as oleic acid (18:1), linoleic acid (18:2), and arachidonic acid (20:4), were nearly undetectable in urine in any patients with various levels of proteinuria and estimated glomerular filtration rates (eGFRs). These findings indicate that nearly all albumin-bound FAs filtered into primary urine are reabsorbed by renal tubules independent of albumin, even in patients with severe kidney diseases.

**Figure 7.**
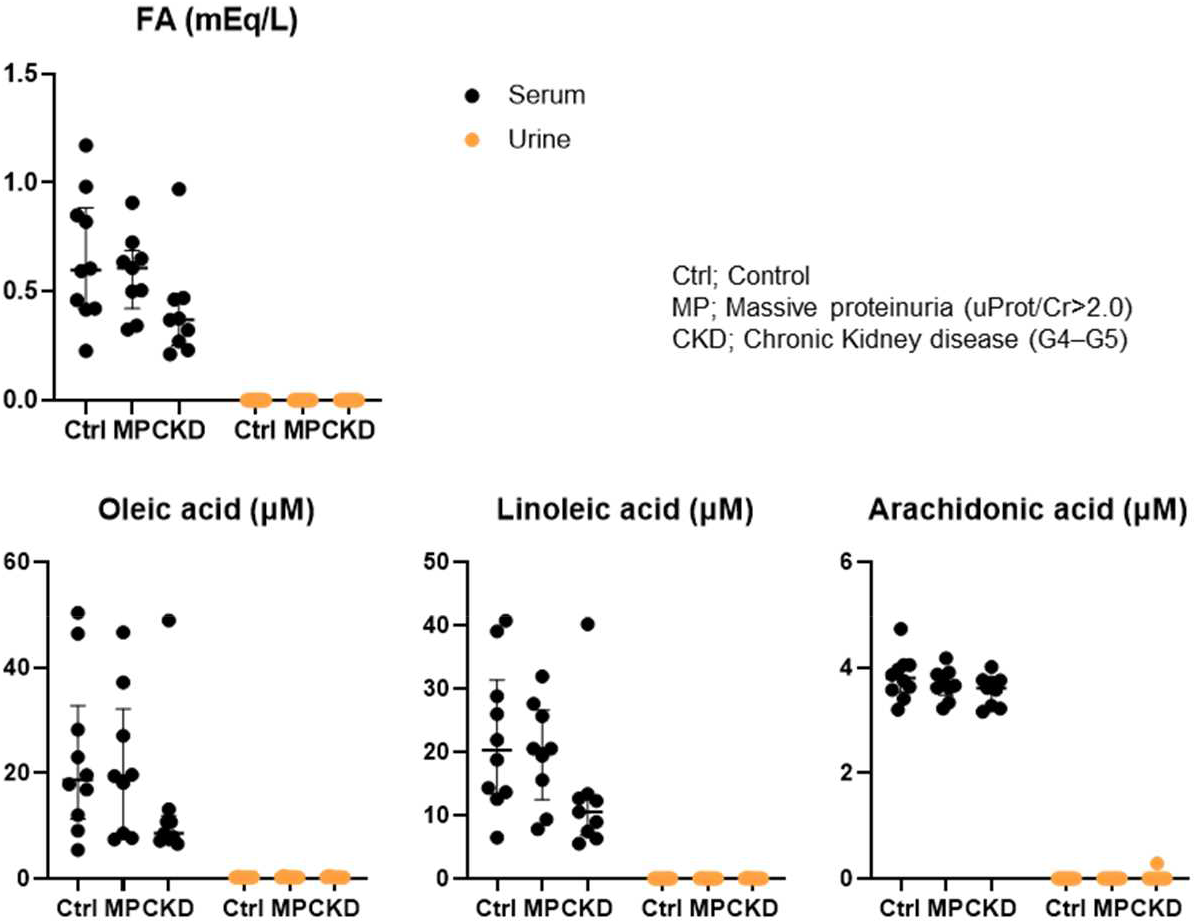
Urine FAs were undetectable even in patients with renal diseases. (Upper panel) Total FA levels in serum and urine were measured with biochemical assays in patients with normal renal function (Ctrl), massive proteinuria (MP), or chronic kidney disease G4−G5 (CKD). (n=9−10). (Lower panel) Oleic acid, linoleic acid, and arachidonic acid levels were measured using liquid chromatography and mass spectrometry.

## Discussion

In the present study, we demonstrated three important findings. First, FAs are not only taken up from the basolateral membrane at least partially via CD36 (or blood) but are also partially reabsorbed from the apical membrane (or primary urine) independent of CD36. FA reabsorption from the apical side was enhanced by the elevation of the serum FA concentration (Fig 8). Second, neutral lipid accumulation occurs from the proximal site of the nephron (S1/S2) when serum FA levels are increased via accelerated lipolysis. In addition, the downstream nephron segments can store excess FAs as TG droplets when PTECs are specifically injured. Third, albumin-bound FAs in the primary urine are completely reabsorbed by the nephron, thus resulting in no detectable FAs in urine, even in acute PTEC injury model mice with overt proteinuria and in humans with massive proteinuria and/or deteriorated kidney function. We discuss each point of these findings and related issues in detail below.

**Figure 8.**
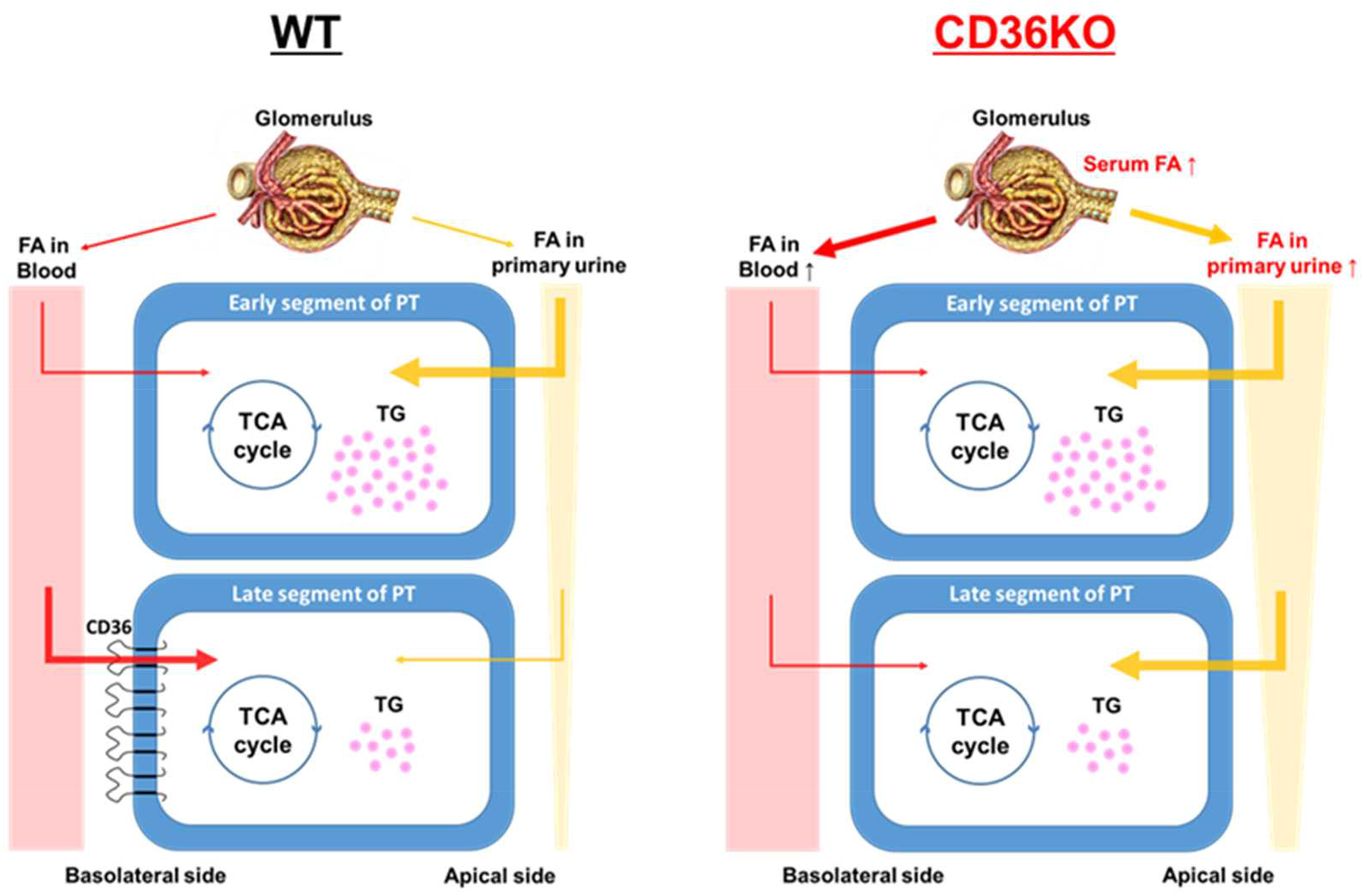
Schematic diagram of bidirectional FA uptake by PTECs in WT and CD36KO mice. In WT mice, the late segments of the proximal tubule (PT) take up FAs from the blood via CD36, while the early segments without CD36 expression take up FAs mainly from primary urine. In CD36KO mice, FA uptake from blood is reduced in the late segments, but the FA concentration in primary mine is elevated, compensating for the reduced FA uptake from the blood. The details are described in the discussions section.

### Bidirectional FA uptake in the kidney

FA uptake from the basolateral membrane (blood) is supported by the following two findings. First, BODIPY-C_12_ was first detected in the peritubular capillary and basolateral membrane in the late segment of PTECs. Second, CD36 is abundantly expressed in the basolateral membrane of PTECs, and ^125^I-BMIPP uptake was reduced in CD36KO mice shortly after its injection. The same distribution pattern as CD36 was also observed in LPL (Nyren et al., 2019). This finding suggests that FAs liberated from lipoproteins after lipolysis also seem to be taken up from the blood. Previous studies have reported that CD36 and LPL are not involved in FA uptake in the kidney (Scerbo et al., 2017), but those data could be due to masking via compensatory FA reabsorption from the primary urine.

FA uptake from the apical membrane (primary urine) is supported by the following four findings. First, BODIPY-C_12_ accumulation was present in the brush border of PTECs, especially in the early segment of the PTs. Second, TGs accumulate even in S3/TAL/DCT when a large amount of TGs is detected in the kidney. TGs also accumulate in S3/TAL/DCT/cortical CD when PTECs are specifically injured. These findings suggest that FAs that are not reabsorbed in S1/S2 are reabsorbed and stored as TGs in the subsequent nephron segments. Third, the reduction in ^125^I-BMIPP uptake in CD36KO kidneys shortly after intravenous injection disappeared in the late phase, which could be due to compensatory reabsorption from the primary urine (Fig 8). Fourth, there were no detectable FAs in the urine in megalin KO and PTi mice, despite prominent proteinuria. Likewise, there were no detectable FAs in the urine in patients with massive proteinuria and/or deteriorated kidney function. These findings suggest that abundant FAs bound to albumin in the primary urine are completely reabsorbed from the apical membrane throughout the entire nephron. Thus, it is very likely that FAs are taken up from both apical and basolateral membranes in the kidney.

### Molecular mechanism of FA reabsorption from primary urine

Previous studies suggested several candidate molecules that might play a role in FA uptake from primary urine. These candidate molecules include megalin, fatty acid transport protein 2 (FATP2), and acyl-CoA synthetase long chain family member 1 (ACSL), whose expression is abundant in PTECs (Christensen et al., 2012, Khan et al., 2014, Khan et al., 2018). In this study, we excluded the possibility of megalin as a candidate to regulate FA reabsorption. Ex vivo studies have reported that FATP2 plays a role in FA absorption from the apical side (Khan et al., 2018). ACSL1 is expressed below the apical membrane and has acyl-CoA synthase activity, which may affect FA uptake from the apical side (Khan et al., 2014). It is warranted to examine whether FATP2 and ACSL1 play a role in FA uptake from primary urine *in vivo*.

### Renal lipid accumulation by elevation of serum FA levels is mediated by increased FA concentration in primary urine but independent of albumin reabsorption

When serum FA levels were increased, lipid droplets appeared in S1/S2, followed by S3, TAL, and DCT. A close association between serum FA levels and TG accumulation has been previously reported (Scerbo et al., 2017). It is also well known that a certain amount of plasma albumin is filtered through the glomerulus, even in intact kidneys (Bobulescu, 2010, Dickson et al., 2014, Comper et al., 2008, Birn and Christensen, 2006), which was confirmed by examining urinary albumin in megalin KO mice in this study. Indeed, BODIPY-C_12_ accumulation was present in the brush border of PTECs, thus strongly suggesting the filtration of albumin-bound FAs into the primary urine. Thus, it is conceivable that FA levels in primary urine are proportional to albumin-bound FA levels in serum. Lipid droplets were also observed in megalin KO mice with defective albumin reabsorption, thus suggesting that FA uptake from the primary urine is independent of megalin and albumin reabsorption. When PTECs were injured by the DT/DT-R system, lipid droplets in the downstream nephron segments (S3/TAL/DCT/cortical CD) were increased, thus suggesting that FA reabsorption in these segments is also dependent on the concentration of FAs in the primary urine but independent of megalin and albumin reabsorption. Moreover, lipid droplets were barely detected in tDL/tAL/medullary CD, even when lipid accumulation reached a maximum value. These segments are believed to not consume FAs as the main energy substrate (Soltoff, 1986, Tian and Liang, 2021). Thus, we suggest that lipid accumulation primarily occurs in FA-dependent segments, such as PTs, TALs, and DCTs, when FA levels are elevated in the primary urine.

### TG accumulation is mainly caused by suppressed FA oxidation and/or excess FA supply in injured kidneys

TG accumulation is increased in both uninjured and injured kidneys. Acute and chronic kidney injuries are thought to cause mitochondrial dysfunction, thus resulting in reduced FA oxidation (Bhargava and Schnellmann, 2017). Modest TG accumulation appeared in the kidney in PTi mice without CL316,243 administration (Figs 6A and 6D), which is presumably due to suppressed FA oxidation. Lipid accumulation in injured segments was enhanced by CL316,243 administration (Fig 6D), thus suggesting TG accumulation via both suppressed FA oxidation and increased FA uptake. Similar TG accumulation is also reported in mouse kidneys with many pathological situations, such as sepsis and ischemia–reperfusion injury (Zager et al., 2005, Tannenbaum et al., 1983, Kang et al., 2015), which further supports the notion that reduced FA oxidation causes lipid accumulation. In acute kidney injury models, it has also been suggested that phospholipase A2, which is an enzyme that hydrolyzes membrane phospholipids to release arachidonic acids, is involved in TG accumulation (to a certain degree) (Zager et al., 2005). Taken together, it is plausible that TG accumulation in injured kidneys is caused by a combination of suppressed FA oxidation in tubular cells, increased FA supply from the blood/primary urine, and FAs derived from injured plasma membranes. It is necessary to determine the contribution of each factor to lipid droplet formation in different pathophysiological situations.

### Is TG accumulation a cause of kidney injury?

There is increasing evidence showing the association between lipid accumulation and renal dysfunction in animal models, as described above. In general, excessive intracellular FAs can be converted into toxic metabolites, such as ceramide and diacylglycerol, under diseased conditions, which is believed to induce cellular damage and mitochondrial dysfunction (Bobulescu, 2010). However, it is still controversial as to whether TG accumulation is a direct cause of kidney injury. There are both pros and cons for the thesis. Pros: Renal dysfunction that is induced by several models, including unilateral ureteral obstruction, diabetes, and the administration of toxic reagents, was alleviated in mice with reduced FA utilization via the genetic deletion of CD36 and FATP2 (Okamura et al., 2009, Yang et al., 2017, Khan et al., 2018, Khan et al., 2020). Likewise, the antagonism of CD36 by the 5A peptide prevented chronic kidney disease progression in mice (Souza et al., 2016). These findings suggest that lipid overload exacerbates kidney injuries induced by other etiologies and can act as a facilitator of kidney injuries.

Cons: Although TG often serves as a measurable indicator of lipid overload, TG is not considered toxic per se. Overnight fasting induces a marked increase in TG accumulation with reduced levels of ceramide (Scerbo et al., 2017). In addition, a high-fat diet alone exerts TG accumulation in the kidney without overt renal dysfunction in mice (Jiang et al., 2005, Yang et al., 2017), which is supported by the rare occurrence of obesity-related glomerulopathy in humans, despite its high prevalence of obesity (Kambham et al., 2001). Furthermore, tubular epithelial cell-specific CD36 overexpression leads to marked lipid accumulation (but few profibrotic changes) in mouse kidneys (Kang et al., 2015). These findings suggest that TG accumulation alone is not harmful to the kidney. In humans and animals, renal fibrosis with TG accumulation is associated with defective FA oxidation (Kang et al., 2015). Indeed, pharmacological interventions for stimulating FA oxidation were found to ameliorate renal fibrosis (Kang et al., 2015). These findings suggest that accumulated TG results from reduced FA oxidation via kidney injury and is not a cause of injury.

The lipotoxicity hypothesis regarding the heart is also under debate (Iso and Kurabayashi, 2021, Ritterhoff et al., 2020, Kenny and Abel, 2019, Jia et al., 2018, Schulze et al., 2016). The modulation of FA uptake in the heart has exhibited both positive and negative effects on cardiac dysfunction in various disease models (Koonen et al., 2007, Yang et al., 2007, Umbarawan et al., 2018b, Umbarawan et al., 2018a, Umbarawan et al., 2020, Umbarawan et al., 2021, Steinbusch et al., 2011, Sung et al., 2017). Thus, further studies are needed to clarify whether accumulated TG is a cause or a result (a beneficial or negative factor) for the development of overnutrition-associated kidney diseases.

### Physiological significance of undetectable FAs in urine

Our data highlight the robust capability of TECs to reabsorb FAs. In humans and animals with massive albuminuria, abundant FAs bound to albumin should also exist in the primary urine. However, to the best of our knowledge, there have been no studies reporting obvious FAs in the urine. This is in significant contrast to readily detectable proteinuria and glucosuria conditions that are observed in kidney diseases and diabetes. Although the term lipiduria exists, this term only means urine containing lipoprotein-rich sediments (but not FA-rich urine) (Blackburn et al., 1998). TGs are the most suitable form for long-term energy storage and supply among various energy substrates. There are many advantages concerning the use of FAs and TGs for systemic energy homeostasis for survival (e.g., a large capacity for TG storage in the body, long-term energy supply, highest energy production per weight, and materials for ketogenesis). Serum FAs are provided from TGs via lipolysis, which is enhanced by adrenergic stimuli, such as fasting, exercise, and cold. Our findings of no detectable FAs in the urine, even in diseased kidneys, suggest that the kidney plays an important role in the regulation of systemic metabolism via complete reabsorption for preventing lipid loss. Together, we propose that the kidney acts not only as a FA-consuming tissue but also as a FA-keeping tissue (or as a thrifty tissue that completely reabsorbs FAs from primary urine) for unwasted energy use against life-threatening conditions, such as starvation and cold.

## Materials and methods

### Animal models

CD36-deficient (CD36KO) mice with a C57BL/6J background were generated as described previously (Putri et al., 2015). Cre-inducible diphtheria toxin receptor (iDTR) mice were purchased from The Jackson Laboratory (Buch et al., 2005) (Bar Harbor, ME). *Ndrg1*^*CreERT2/+*^ mice (Endo et al., 2015) were crossed with iDTR mice to generate *Ndrg1*^*CreERT2/+*^*:iDTR* mice. The proximal tubule injury (PTi) model was produced as described previously (Takaori et al., 2016). In brief, 0.15 mg/g body weight tamoxifen (T5648, Sigma– Aldrich, St. Louis, MO) was intraperitoneally administered to *Ndrg1*^*CreERT2/+*^*:iDTR* mice for 5 consecutive days (Higashi et al., 2009). Three to four weeks later, 25 ng/g body weight diphtheria toxin (Sigma–Aldrich, St. Louis, MO) was intraperitoneally administered to the mice. PT-specific megalin KO mice (*Ndrg1*^*CreERT2/+*^*:megalin*^*fl/fl*^) with the C57BL/6J background were generated as described elsewhere (Kuwahara et al., 2016). To delete the megalin gene in adulthood, 175 mg/kg body weight tamoxifen dissolved in sunflower oil/ethanol (11:1) (17.5 mg/mL) was gavaged to mice for 5 days a week at the age of 7 and 9 weeks. These mice were housed in a temperature-controlled room (20–26 °C) with a 12-h light/12-h dark cycle and given unrestricted access to water and standard chow (CE-2, Clea Japan, Inc.). Littermates were used as controls for all studies. Mice were euthanized using 2% isoflurane and cervical dislocation. Male mice aged 7–12 weeks were used to avoid estrogen cycle-related confounding effects. All mouse strains used in this study were backcrossed to C57BL/6J mice for at least three generations.

To induce lipolysis, 3 adrenergic receptor agonists, CL316,243 (TOCRIS No. 1499), were intraperitoneally injected into mice at a dose of 1 mg/kg.

### Histological study and immunostaining

The kidneys were fixed with 4% paraformaldehyde or Carnoy solution (ethanol: chloroform: acetic acid = 6:3:1) and embedded in paraffin. Immunohistochemistry (IHC) or immunofluorescence (IF) was performed with the following antibodies and lectin: CD36 (Abcam, ab124515, rabbit polyclonal); Aquaporin1 (AQP1), AQP2, and AQP4 (affinity-purified rabbit antibody (Matsuzaki et al., 2009)); Na^+^-Cl^-^ cotransporter (NCC) (Chemicon, AB3553, rabbit polyclonal); Tamm-Horsfall protein (THP) (Santa Cruz, sc-19554, goat polyclonal); and Lotus Tetragonolobus Lectin (LTL) Fluorescein (Vector, FL-1321). Nuclei were stained with hematoxylin for IHC or 4′,6-diamidino-2-phenylindole (DAPI) for IF.

To assess neutral lipid accumulation, fresh unfixed kidneys were embedded in Tissue-Tek OCT compound (SAKURA, 83-1824, Japan) and sliced using a CM3050S cryostat (Leica, Germany). Eight-micrometer cryosections fixed with 4% paraformaldehyde for 10 min were stained with oil-red O. To identify lipid-accumulating cells, simultaneous fluorescent analysis was performed with LTL fluorescein and several antibodies to detect the nephron structure.

All images were obtained with a BZ-9000 fluorescence microscope (Keyence, Osaka, Japan)

### *In vivo* imaging of BODIPY-C12 FAs with multiphoton microscopy

*In vivo* imaging with multiphoton microscopy was performed as described previously (Nakano et al., 2015). A fluorescence-labeled medium-chain FA tracer, 4,4-difluoro-5-(2-thienyl)-4-bora-3a,4a-diaza-s-indacene-3-dodecanoic acid (BODIPY-C_12_, Invitrogen D3835), was dissolved in 100% ethanol and conjugated to 10% bovine serum albumin. After an overnight fast, BODIPY-C_12_ was injected intravenously via the cervical vein at a dose of 150 nmol/mouse, and *in vivo* imaging was performed for 30 min using an Olympus FV1000MPE multiphoton confocal fluorescence imaging system (Olympus, Tokyo, Japan).

### Lipid extraction and measurement

Lipids were extracted from tissues with the modified Folch method (Folch et al., 1957). In brief, 40 mg of tissue was homogenized in 800 μl of chloroform/methanol (2:1) and centrifuged for 15 min at 12,000 g at room temperature. One hundred forty microliters of distilled water were added to 700 μl of the supernatant and vortexed. After centrifugal separation for 20 min at 18,000 g at room temperature, the lower organic phase was collected and blown dry with nitrogen gas. The dried lipid was dissolved with 50 μl of isopropanol. TGs (Triglyceride E-test, Wako Chemical, Osaka, Japan) and FAs (NEFA C-test, Wako Chemical) were measured according to the manufacturer’s protocols.

### Biodistribution of ^125^I-BMIPP (15-(p-iodophenyl)-3-(R,S)-methyl pentadecanoic acid) and ^18^F-FDG (2-fluorodeoxyglucose)

The biodistribution of ^125^I-BMIPP and ^18^F-FDG was determined as described previously (Coburn et al., 2000, Hajri et al., 2002). Mice received intravenous injections of ^125^I-BMIPP (5 kBq) and ^18^F-FDG (100 kBq) via the lateral tail vein in a volume of 100 μL. ^125^I-BMIPP was a gift from Nihon Medi-Physics Co. Ltd., and ^18^F-FDG was obtained from batches prepared for clinical PET imaging at Gunma University. The isolated tissues were weighed and counted in a well-type gamma counter (ARC-7001, ALOKA). When blood removal was necessary, mice were perfused with 10 mL PBS before tissue isolation. The data are expressed as % injected dose/gram (%ID/g). Autoradiography for ^125^I-BMIPP was performed with a 1 mm slice of unfixed kidney and an imaging plate (BAS-MS2025, Fujifilm, Tokyo, Japan).

### Human samples

Twenty-eight serum and urine samples that were collected at Gunma University Hospital between October 2011 and March 2018 were used for lipid measurements. They included samples from 9 patients with massive proteinuria (urine protein/creatinine > 2.0 g/gCr), 9 with chronic kidney disease (CKD) glomerular filtration rate (GFR) categories 4 or 5 (G4–G5: GFR < 29 mL/min/1.73 mm^2^), and 10 with normal kidney function (GFR > 60 mL/min/1.73 mm^2^ without hyperproteinuria). FAs (NEFA-HR, Wako Chemical), TGs (Triglyceride kit L-type TG-M, Wako Chemical), and total cholesterol (Cholesterol kit L-type CHO-M, Wako Chemical) were measured according to the manufacturer’s protocols. Nonesterified fatty acid species were measured using a triple quadrupole mass spectrometer coupled with a liquid chromatography (LCMS-8050 system, Shimadzu). Fatty acids were extracted from serum and urine in chloroform/methanol (2/1). After centrifugation, the lower organic phase was collected and dried with nitrogen gas and then resuspended in 80% methanol. The extract was separated on a reversed-phase C18 column (Mastro C18, 2.1 mm × 150 mm, 3 μm, Shimadzu) by using a gradient of solvents A (10 mM ammonium acetate in water) and B (acetonitrile) with a flow rate of 0.25 ml/min. The initial solvent composition was 80% B, and the following solvent gradient was applied: 80% B for 2 min, increased linearly to 98% from 2 to 12 min, held at 98% B for 5 min, then returned to 80% B and maintained for 5 min. The column was maintained at 40 °C.

The separated analytes were ionized by electrospray ionization and then measured by mass spectrometry in selected reaction monitoring (SRM) mode. The SRM transitions for the analytes were *m/z* 281.2 > 281.2 [M –H]^−^ for oleic acid, *m/z* 279.2 > 279.2 [M –H]^−^ for linoleic acid, and *m/z* 303.2 > 303.2 [M –H]^−^ for arachidonic acid. The peak height of each fatty acid was applied to the standard curve made for each fatty acid species for quantification.

### Statistical analysis

Statistical analysis was performed with GraphPad Prism 8 software. The data are presented as the mean ± standard deviation. Student’s t test was performed for 2-group comparisons. One-way analysis of variance (ANOVA) with Tukey *post hoc* multiple comparison tests was carried out for 3-group comparisons. Two-way ANOVA was used to analyze the effects of genotype (Control vs. CD36KO, megalin KO, PTi mice), treatments (fasted, refed or CL316,243), and their interaction. A p value < 0.05 was considered to be statistically significant. *p<0.05, **p<0.01, ***p<0.001: main effect for genotype. ^#^p<0.05, ^##^p<0.01, ^###^p<0.001: main effect for treatments. ^†^p<0.05, ^††^p<0.01, ^†††^p<0.001: interaction between genotype and treatments.

### Study approval

All animal studies were approved by the Institutional Animal Care and Use Committee (Gunma University Graduate School of Medicine, approval number: 19-014 and 20-055) and conformed to the NIH guidelines (Guide for the Care and Use of Laboratory Animals). The human study was approved by the Institutional Review Board at Gunma University Hospital (approval number: HS2021-060). Written informed consent was received from all patients prior to participation.

## Acknowledgments

We thank Takako Kobayashi and Keiko Matsukura for their excellent technical assistance. This work was supported in part by Grants-in-Aid for Scientific Research from the Japan Society for the Promotion of Science to MK (20H03671) and TI (20K08878); grants from Japan Agency for Medical Research and Development to MK (20gm0910003h0106); AS (JP20am0101094) and MY (21gm1210009, 21zf0127003h001); and a grant from the Naito Foundation to TI.

## Author contributions

Conceptualization: TI, Data curation: RK, HI, KH, Formal analysis: RK, HH, HO, DN, Funding acquisition: MK, TI, AS, MY, Investigation: RK, HH, AK, HO, DN, MM, RT, HS, SG, HM, NK, KS, TY, HI, TI, Methodology: HH, AK, HO, DN, AN, Project Administration: HI, KH, TI, Resources: HI, TM, KH, AS, MY, Supervision: MK, Visualization: RK, TI, Writing – original draft, RK, TI, Writing – review & editing: RK, HO, DN, HI, KH, MY, MK, TI.

## Data and material availability

All data are available in the main text and the supplementary materials.

## References

Abumrad, N. A., Cabodevilla, A. G., Samovski, D., Pietka, T., Basu, D. & Goldberg, I. J. 2021. Endothelial Cell Receptors in Tissue Lipid Uptake and Metabolism. Circulation Research, 128, 433–450. doi:10.1161/CIRCRESAHA.120.318003

Bhargava, P. & Schnellmann, R. G. 2017. Mitochondrial energetics in the kidney. Nat Rev Nephrol, 13, 629–646. 10.1038/nrneph.2017.107

Birn, H. & Christensen, E. I. 2006. Renal albumin absorption in physiology and pathology. Kidney Int, 69, 440–9. 10.1038/sj.ki.5000141

Blackburn, V., Grignani, S. & Fogazzi, G. B. 1998. Lipiduria as seen by transmission electron microscopy. Nephrology Dialysis Transplantation, 13, 2682–2684. 10.1093/ndt/13.10.2682

Bobulescu, I. A. 2010. Renal lipid metabolism and lipotoxicity. Curr Opin Nephrol Hypertens, 19, 393–402. 10.1097/MNH.0b013e32833aa4ac

Buch, T., Heppner, F. L., Tertilt, C., Heinen, T. J., Kremer, M., Wunderlich, F. T., Jung, S. & Waisman, A. 2005. A Cre-inducible diphtheria toxin receptor mediates cell lineage ablation after toxin administration. Nat Methods, 2, 419–26. 10.1038/nmeth762

Christensen, E. I., Birn, H., Storm, T., Weyer, K. & Nielsen, R. 2012. Endocytic Receptors in the Renal Proximal Tubule. Physiology, 27, 223–236. 10.1152/physiol.00022.2012

Coburn, C. T., Knapp, F. F., JR., Febbraio, M., Beets, A. L., Silverstein, R. L. & Abumrad, N. A. 2000. Defective uptake and utilization of long chain fatty acids in muscle and adipose tissues of CD36 knockout mice. J Biol Chem, 275, 32523–9. 10.1074/jbc.M003826200

Comper, W. D., Haraldsson, B. & Deen, W. M. 2008. Resolved: normal glomeruli filter nephrotic levels of albumin. J Am Soc Nephrol, 19, 427–32. 10.1681/asn.2007090997

Dickson, L. E., Wagner, M. C., Sandoval, R. M. & Molitoris, B. A. 2014. The proximal tubule and albuminuria: really! J Am Soc Nephrol, 25, 443–53. 10.1681/ASN.2013090950

Endo, T., Nakamura, J., Sato, Y., Asada, M., Yamada, R., Takase, M., Takaori, K., Oguchi, A., Iguchi, T., Higashi, A. Y., Ohbayashi, T., Nakamura, T., Muso, E., Kimura, T. & Yanagita, M. 2015. Exploring the origin and limitations of kidney regeneration. J Pathol, 236, 251–63. 10.1002/path.4514

Folch, J., Lees, M. & Sloane Stanley, G. H. 1957. A simple method for the isolation and purification of total lipides from animal tissues. J Biol Chem, 226, 497–509.

Gallagher, D., Belmonte, D., Deurenberg, P., Wang, Z., Krasnow, N., Pi-Sunyer, F. X. & Heymsfield, S. B. 1998. Organ-tissue mass measurement allows modeling of REE and metabolically active tissue mass. Am J Physiol, 275, E249–58. 10.1152/ajpendo.1998.275.2.E249

Hajri, T., Han, X. X., Bonen, A. & Abumrad, N. A. 2002. Defective fatty acid uptake modulates insulin responsiveness and metabolic responses to diet in CD36-null mice. J Clin Invest, 109, 1381–9. 10.1172/JCI14596

Hasan, S. S. & Fischer, A. 2021. The Endothelium: An Active Regulator of Lipid and Glucose Homeostasis. Trends Cell Biol, 31, 37–49. 10.1016/j.tcb.2020.10.003

Higashi, A. Y., Ikawa, T., Muramatsu, M., Economides, A. N., Niwa, A., Okuda, T., Murphy, A. J., Rojas, J., Heike, T., Nakahata, T., Kawamoto, H., Kita, T. & Yanagita, M. 2009. Direct hematological toxicity and illegitimate chromosomal recombination caused by the systemic activation of CreERT2. J Immunol, 182, 5633–40. 10.4049/jimmunol.0802413

Iso, T. & Kurabayashi, M. 2017. Fatty Acid Uptake by the Heart During Fasting. In: Preedy, V. & Patel, V. B. (eds.) Handbook of Famine, Starvation, and Nutrient Deprivation: From Biology to Policy. Cham: Springer International Publishing.

Iso, T. & Kurabayashi, M. 2021. Cardiac Metabolism and Contractile Function in Mice with Reduced Trans-Endothelial Fatty Acid Transport. Metabolites, 11, 889.

Jia, G., Hill, M. A. & Sowers, J. R. 2018. Diabetic Cardiomyopathy: An Update of Mechanisms Contributing to This Clinical Entity. Circ Res, 122, 624–638. 10.1161/circresaha.117.311586

Jiang, T., Wang, Z., Proctor, G., Moskowitz, S., Liebman, S. E., Rogers, T., Lucia, M. S., Li, J. & Levi, M. 2005. Diet-induced Obesity in C57BL/6J Mice Causes Increased Renal Lipid Accumulation and Glomerulosclerosis via a Sterol Regulatory Element-binding Protein-1c-dependent Pathway*. Journal of Biological Chemistry, 280, 32317-32325. https://doi.org/10.1074/jbc.M500801200

Kambham, N., Markowitz, G. S., Valeri, A. M., Lin, J. & D’Agati, V. D. 2001. Obesity-related glomerulopathy: an emerging epidemic. Kidney Int, 59, 1498–509. 10.1046/j.1523-1755.2001.0590041498.x

Kang, H. M., Ahn, S. H., Choi, P., Ko, Y. A., Han, S. H., Chinga, F., Park, A. S., Tao, J., Sharma, K., Pullman, J., Bottinger, E. P., Goldberg, I. J. & Susztak, K. 2015. Defective fatty acid oxidation in renal tubular epithelial cells has a key role in kidney fibrosis development. Nat Med, 21, 37–46. 10.1038/nm.3762

Kenny, H. C. & Abel, E. D. 2019. Heart Failure in Type 2 Diabetes Mellitus. Circ Res, 124, 121–141. 10.1161/circresaha.118.311371

Khan, S., Abu Jawdeh, B. G., Goel, M., Schilling, W. P., Parker, M. D., Puchowicz, M. A., Yadav, S. P., Harris, R. C., El-Meanawy, A., Hoshi, M., Shinlapawittayatorn, K., Deschenes, I., Ficker, E. & Schelling, J. R. 2014. Lipotoxic disruption of NHE1 interaction with PI(4,5)P2 expedites proximal tubule apoptosis. J Clin Invest, 124, 1057–68. 10.1172/JCI71863

Khan, S., Cabral, P. D., Schilling, W. P., Schmidt, Z. W., Uddin, A. N., Gingras, A., Madhavan, S. M., Garvin, J. L. & Schelling, J. R. 2018. Kidney Proximal Tubule Lipoapoptosis Is Regulated by Fatty Acid Transporter-2 (FATP2). J Am Soc Nephrol, 29, 81–91. 10.1681/ASN.2017030314

Khan, S., Gaivin, R., Abramovich, C., Boylan, M., Calles, J. & Schelling, J. R. 2020. Fatty acid transport protein-2 regulates glycemic control and diabetic kidney disease progression. JCI Insight, 5. 10.1172/jci.insight.136845

Koonen, D. P., Febbraio, M., Bonnet, S., Nagendran, J., Young, M. E., Michelakis, E. D. & Dyck, J. R. 2007. CD36 expression contributes to age-induced cardiomyopathy in mice. Circulation, 116, 2139–47. 10.1161/circulationaha.107.712901

Kuwahara, S., Hosojima, M., Kaneko, R., Aoki, H., Nakano, D., Sasagawa, T., Kabasawa, H., Kaseda, R., Yasukawa, R., Ishikawa, T., Suzuki, A., Sato, H., Kageyama, S., Tanaka, T., Kitamura, N., Narita, I., Komatsu, M., Nishiyama, A. & Saito, A. 2016. Megalin-Mediated Tubuloglomerular Alterations in High-Fat Diet–Induced Kidney Disease. Journal of the American Society of Nephrology, 27, 1996–2008. 10.1681/asn.2015020190

Matsuzaki, T., Hata, H., Ozawa, H. & Takata, K. 2009. Immunohistochemical localization of the aquaporins AQP1, AQP3, AQP4, and AQP5 in the mouse respiratory system. Acta Histochem Cytochem, 42, 159–69. 10.1267/ahc.09023

Mount, P., Davies, M., Choy, S. W., Cook, N. & Power, D. 2015. Obesity-Related Chronic Kidney Disease-The Role of Lipid Metabolism. Metabolites, 5, 720–32. 10.3390/metabo5040720

Nakano, D., Doi, K., Kitamura, H., Kuwabara, T., Mori, K., Mukoyama, M. & Nishiyama, A. 2015. Reduction of Tubular Flow Rate as a Mechanism of Oliguria in the Early Phase of Endotoxemia Revealed by Intravital Imaging. J Am Soc Nephrol, 26, 3035–44. 10.1681/ASN.2014060577

Nyren, R., Makoveichuk, E., Malla, S., Kersten, S., Nilsson, S. K., Ericsson, M. & Olivecrona, G. 2019. Lipoprotein lipase in mouse kidney: effects of nutritional status and high-fat diet. Am J Physiol Renal Physiol, 316, F558–F571. 10.1152/ajprenal.00474.2018

Okamura, D. M., Pennathur, S., Pasichnyk, K., Lopez-Guisa, J. M., Collins, S., Febbraio, M., Heinecke, J. & Eddy, A. A. 2009. CD36 regulates oxidative stress and inflammation in hypercholesterolemic CKD. J Am Soc Nephrol, 20, 495–505. 10.1681/ASN.2008010009

Pi, X., Xie, L. & Patterson, C. 2018. Emerging Roles of Vascular Endothelium in Metabolic Homeostasis. Circ Res, 123, 477–494. 10.1161/circresaha.118.313237

Putri, M., Syamsunarno, M. R., Iso, T., Yamaguchi, A., Hanaoka, H., Sunaga, H., Koitabashi, N., Matsui, H., Yamazaki, C., Kameo, S., Tsushima, Y., Yokoyama, T., Koyama, H., Abumrad, N. A. & Kurabayashi, M. 2015. CD36 is indispensable for thermogenesis under conditions of fasting and cold stress. Biochem Biophys Res Commun, 457, 520–5. 10.1016/j.bbrc.2014.12.124

Ritterhoff, J., Young, S., Villet, O., Shao, D., Neto, F. C., Bettcher, L. F., Hsu, Y.-W. A., Kolwicz, S. C., Raftery, D. & Tian, R. 2020. Metabolic Remodeling Promotes Cardiac Hypertrophy by Directing Glucose to Aspartate Biosynthesis. Circulation Research, 126, 182–196. doi:10.1161/CIRCRESAHA.119.315483

Scerbo, D., Son, N. H., Sirwi, A., Zeng, L., Sas, K. M., Cifarelli, V., Schoiswohl, G., Huggins, L. A., Gumaste, N., Hu, Y., Pennathur, S., Abumrad, N. A., Kershaw, E. E., Hussain, M. M., Susztak, K. & Goldberg, I. J. 2017. Kidney triglyceride accumulation in the fasted mouse is dependent upon serum free fatty acids. J Lipid Res, 58, 1132–1142. 10.1194/jlr.M074427

Schulze, P. C., Drosatos, K. & Goldberg, I. J. 2016. Lipid Use and Misuse by the Heart. Circ Res, 118, 1736–51. 10.1161/circresaha.116.306842

Soltoff, S. P. 1986. ATP and the regulation of renal cell function. Annu Rev Physiol, 48, 9–31. 10.1146/annurev.ph.48.030186.000301

Souza, A. C., Bocharov, A. V., Baranova, I. N., Vishnyakova, T. G., Huang, Y. G., Wilkins, K. J., Hu, X., Street, J. M., Alvarez-Prats, A., Mullick, A. E., Patterson, A. P., Remaley, A. T., Eggerman, T. L., Yuen, P. S. & Star, R. A. 2016. Antagonism of scavenger receptor CD36 by 5A peptide prevents chronic kidney disease progression in mice independent of blood pressure regulation. Kidney Int, 89, 809–22. 10.1016/j.kint.2015.12.043

Steinbusch, L. K., Luiken, J. J., Vlasblom, R., Chabowski, A., Hoebers, N. T., Coumans, W. A., Vroegrijk, I. O., Voshol, P. J., Ouwens, D. M., Glatz, J. F. & Diamant, M. 2011. Absence of fatty acid transporter CD36 protects against Western-type diet-related cardiac dysfunction following pressure overload in mice. Am J Physiol Endocrinol Metab, 301, E618–27. 10.1152/ajpendo.00106.2011

Sung, M. M., Byrne, N. J., Kim, T. T., Levasseur, J., Masson, G., Boisvenue, J. J., Febbraio, M. & Dyck, J. R. B. 2017. Cardiomyocyte-specific ablation of CD36 accelerates the progression from compensated cardiac hypertrophy to heart failure. American Journal of Physiology-Heart and Circulatory Physiology, 312, H552–H560. 10.1152/ajpheart.00626.2016

Takaori, K., Nakamura, J., Yamamoto, S., Nakata, H., Sato, Y., Takase, M., Nameta, M., Yamamoto, T., Economides, A. N., Kohno, K., Haga, H., Sharma, K. & Yanagita, M. 2016. Severity and Frequency of Proximal Tubule Injury Determines Renal Prognosis. J Am Soc Nephrol, 27, 2393–406. 10.1681/ASN.2015060647

Tannenbaum, J., Purkerson, M. L. & Klahr, S. 1983. Effect of unilateral ureteral obstruction on metabolism of renal lipids in the rat. American Journal of Physiology-Renal Physiology, 245, F254–F262. 10.1152/ajprenal.1983.245.2.F254

Tian, Z. & Liang, M. 2021. Renal metabolism and hypertension. Nature Communications, 12, 963. 10.1038/s41467-021-21301-5

Trent, C. M., Yu, S., Hu, Y., Skoller, N., Huggins, L. A., Homma, S. & Goldberg, I. J. 2014. Lipoprotein lipase activity is required for cardiac lipid droplet production. J Lipid Res, 55, 645–58. 10.1194/jlr.M043471

Umbarawan, Y., Kawakami, R., Syamsunarno, M. R. A. A., Koitabashi, N., Obinata, H., Yamaguchi, A., Hanaoka, H., Hishiki, T., Hayakawa, N., Sunaga, H., Matsui, H., Kurabayashi, M. & Iso, T. 2020. Reduced fatty acid uptake aggravates cardiac contractile dysfunction in streptozotocin-induced diabetic cardiomyopathy. Scientific Reports, 10, 20809. 10.1038/s41598-020-77895-1

Umbarawan, Y., Kawakami, R., Syamsunarno, M. R. A. A., Obinata, H., Yamaguchi, A., Hanaoka, H., Hishiki, T., Hayakawa, N., Koitabashi, N., Sunaga, H., Matsui, H., Kurabayashi, M. & Iso, T. 2021. Reduced Fatty Acid Use from CD36 Deficiency Deteriorates Streptozotocin-Induced Diabetic Cardiomyopathy in Mice. Metabolites, 11, 881.

Umbarawan, Y., Syamsunarno, M. R. A. A., Koitabashi, N., Obinata, H., Yamaguchi, A., Hanaoka, H., Hishiki, T., Hayakawa, N., Sano, M., Sunaga, H., Matsui, H., Tsushima, Y., Suematsu, M., Kurabayashi, M. & Iso, T. 2018a. Myocardial fatty acid uptake through CD36 is indispensable for sufficient bioenergetic metabolism to prevent progression of pressure overload-induced heart failure. Scientific Reports, 8, 12035. 10.1038/s41598-018-30616-1

Umbarawan, Y., Syamsunarno, M. R. A. A., Koitabashi, N., Yamaguchi, A., Hanaoka, H., Hishiki, T., Nagahata-Naito, Y., Obinata, H., Sano, M., Sunaga, H., Matsui, H., Tsushima, Y., Suematsu, M., Kurabayashi, M. & Iso, T. 2018b. Glucose is preferentially utilized for biomass synthesis in pressure-overloaded hearts: evidence from fatty acid-binding protein-4 and -5 knockout mice. Cardiovascular Research, 114, 1132–1144. 10.1093/cvr/cvy063

Wahl, P., Ducasa, G. M. & Fornoni, A. 2016. Systemic and renal lipids in kidney disease development and progression. Am J Physiol Renal Physiol, 310, F433–45. 10.1152/ajprenal.00375.2015

Weyer, K., Storm, T., Shan, J., Vainio, S., Kozyraki, R., Verroust, P. J., Christensen, E. I. & Nielsen, R. 2011. Mouse model of proximal tubule endocytic dysfunction. Nephrol Dial Transplant, 26, 3446–51. 10.1093/ndt/gfr525

Yang, J., Sambandam, N., Han, X., Gross, R. W., Courtois, M., Kovacs, A., Febbraio, M., Finck, B. N. & Kelly, D. P. 2007. CD36 deficiency rescues lipotoxic cardiomyopathy. Circ Res, 100, 1208–17. 10.1161/01.RES.0000264104.25265.b6

Yang, P., Xiao, Y., Luo, X., Zhao, Y., Zhao, L., Wang, Y., Wu, T., Wei, L. & Chen, Y. 2017. Inflammatory stress promotes the development of obesity-related chronic kidney disease via CD36 in mice. J Lipid Res, 58, 1417–1427. 10.1194/jlr.M076216

Zager, R. A., Johnson, A. C. M. & Hanson, S. Y. 2005. Renal tubular triglyercide accumulation following endotoxic, toxic, and ischemic injury. Kidney International, 67, 111-121. https://doi.org/10.1111/j.1523-1755.2005.00061.x

