## Supplemental data for "Robust capability of renal tubule fatty acid uptake from apical and basolateral membranes in physiology and disease"

Ryo Kawakami^1^, Hirofumi Hanaoka^2^, Ayaka Kanai^2^, Hideru Obinata^3^, Daisuke Nakano^4^, Hidekazu Ikeuchi^5^, Miki Matsui^1^, Toshiyuki Matsuzaki^6^, Rina Tanaka^7^, Hiroaki Sunaga^1,12^, Sawako Goto^8^, Hiroki Matsui^7^, Norimichi Koitabashi^1^, Keiko Saegusa^9^, Tomoyuki Yokoyama^7^, Keiju Hiromura^5^, Akira Nishiyama^4^, Akihiko Saito^8^, Motoko Yanagita^10,11^, Hideki Ishii^1^, Masahiko Kurabayashi^1^, Tatsuya Iso^1,9*^

^1^Department of Cardiovascular Medicine, ^2^Department of Bioimaging Information Analysis, ^3^Education and Research Support Center, ^5^Department of Nephrology and Rheumatology, and ^6^Department of Anatomy and Cell Biology, Gunma University Graduate School of Medicine, Maebashi, Gunma, Japan.

^4^Department of Pharmacology, Faculty of Medicine, Kagawa University, Kitagun, Kagawa, Japan.

^7^Department of Laboratory Sciences, Gunma University Graduate School of Health Sciences, Maebashi, Gunma, Japan.

^8^Department of Applied Molecular Medicine, Kidney Research Center, Niigata University Graduate School of Medical and Dental Sciences, Chuo-ku, Niigata, Japan.

^9^Department of Medical Technology and Clinical Engineering, Faculty of Medical Technology and Clinical Engineering, Gunma University of Health and Welfare, Maebashi, Gunma, Japan.

^10^Department of Nephrology, Graduate School of Medicine, ^11^Institute for the Advanced Study of Human Biology (ASHBi), Kyoto University, Sakyo-ku, Kyoto, Japan.

^12^Center for Liberal Arts and Sciences, Ashikaga University, Ashikaga, Tochigi, Japan.

* Corresponding author

 (TI)

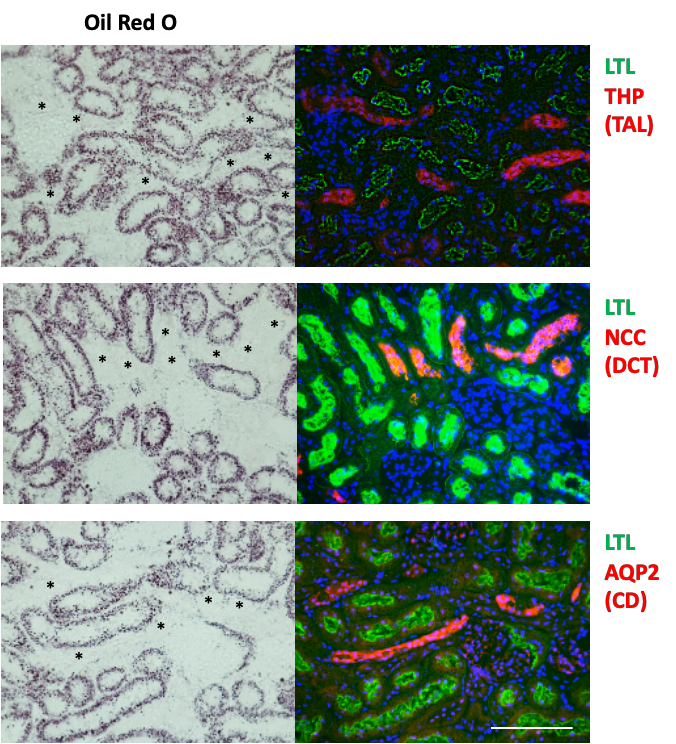

100 µm

**Figure S1. Spatial distribution of the modestly accumulated lipid after CL 316, 243 injections.** Oil-red O staining and immunofluorescence (IF) were simultaneously executed. Representative images showing LTL (green), THP (red), NCC (red), and AQP2 (red). Black asterisks represent IF-positive tubules. Scale bars, 100 µm.

100 µm

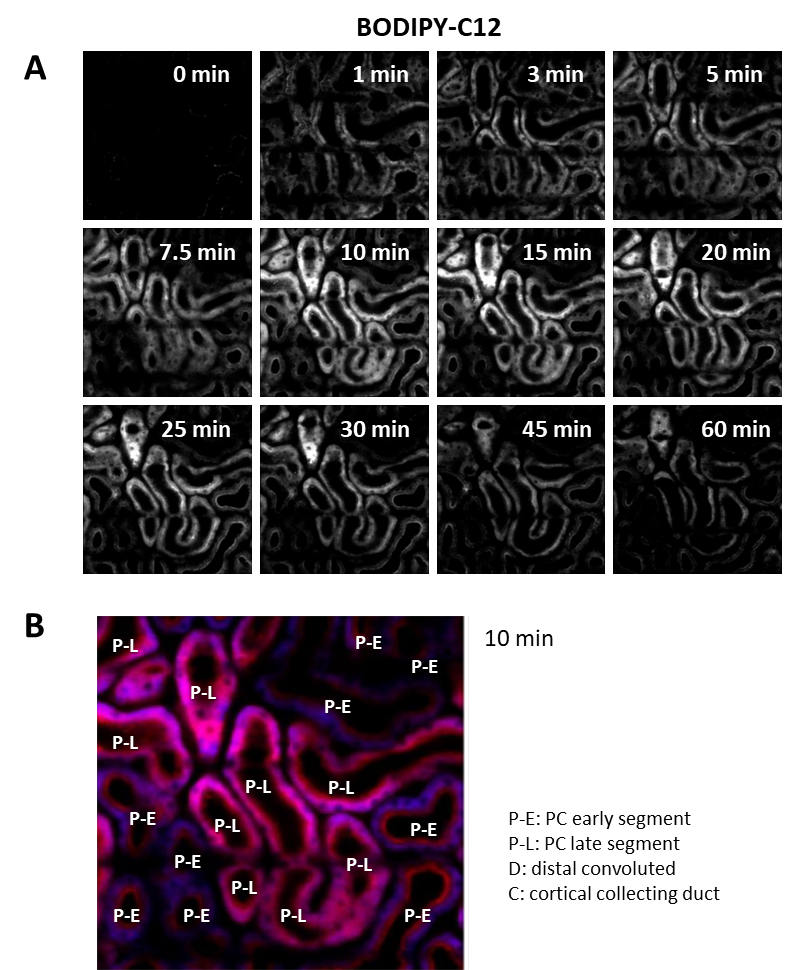

**Figure S2. Time course of BODIPY-C_12_ accumulation in PTECs.** (A) Time-lapse images (0–60 min) with multiphoton microscopy showing 4,4-Difluoro-5-(2-Thienyl)-4-Bora-3a,4a-Diaza-s-Indacene-3-Dodecanoic Acid (BODPY-C_12_) in grey scale. Scale bars, 100µm. (B) The early or late segments of proximal tubules (P-E or P-L, respectively) were indicated on the image at 10 min.

100 µm

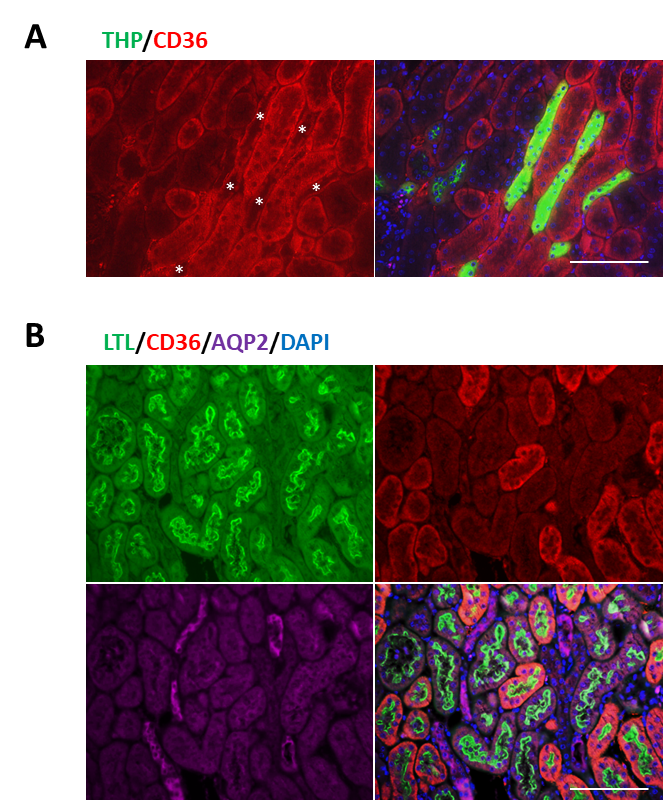

100 µm

100 µm

**Figure S3.** **The distribution of CD36 in mice kidney.** Representative images of IF showing (A) THP (green) and CD36 (red); (B) LTL (green), CD36 (red), AQP2 (purple), and DAPI (blue) in WT kidney. Scale bars, 100 µm.

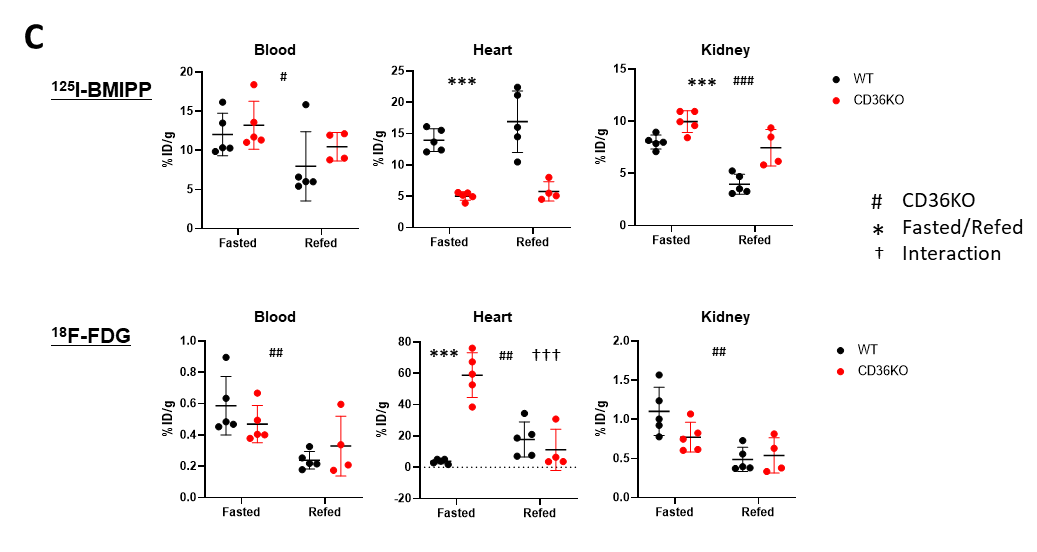

**A**

**C**

**B**

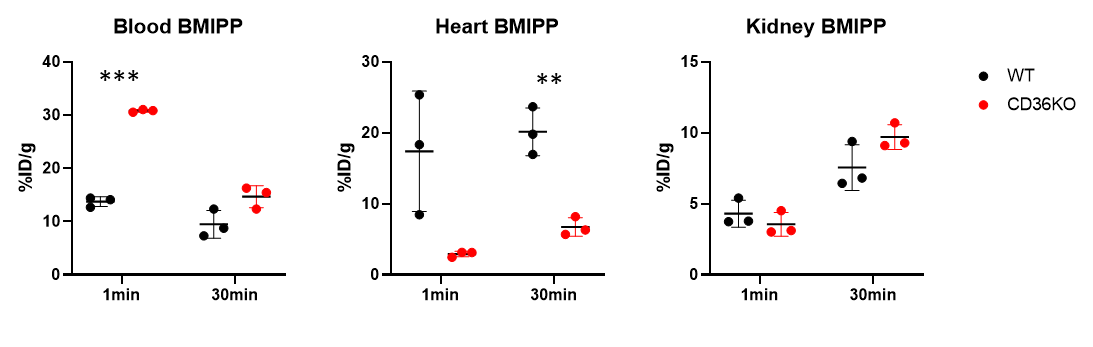

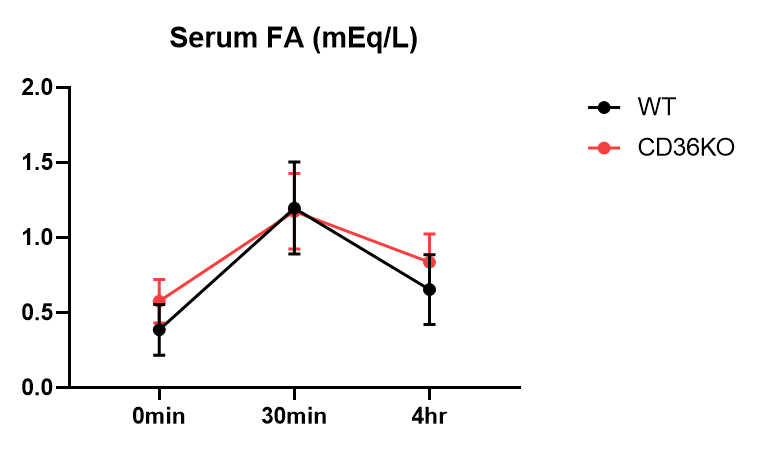

**Figure S4. FA uptake study with ^125^I-BMIPP.** (A) After overnight-fasting with or without refeeding, ^125^I-BMIPP and ^18^F-FDG were injected into the mice. Indicated organs were collected 2 h after the injections. (n = 4–5) (B) Blood was drawn from the retro-orbital plexus 1 or 30 min after ^125^I-BMIPP injections. (n=3) (C) Time course of serum levels of FA after CL 316, 243 injections in WT and CD36KO mice. (n = 6)

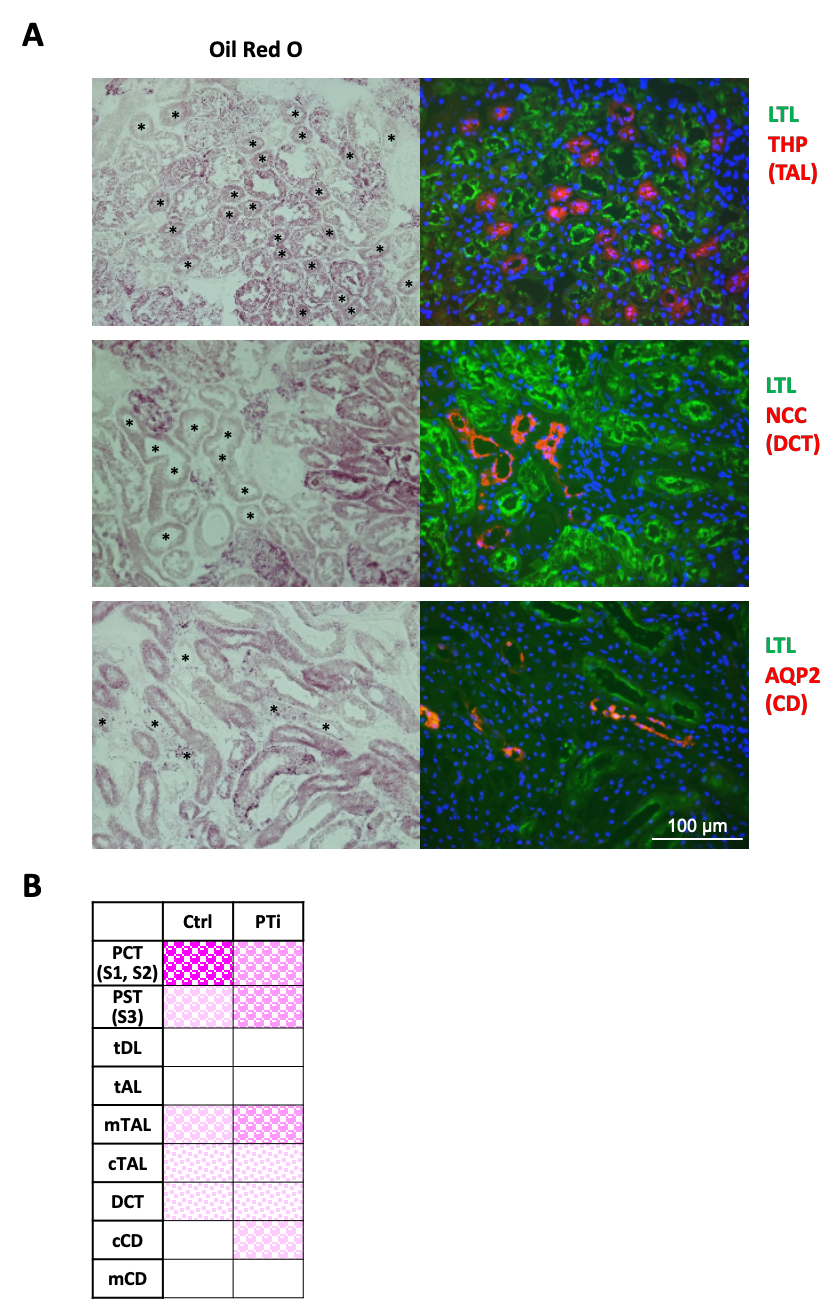

**Figure S5. Lipid distribution after CL 316, 243 injections in PTi mice.** (A) Oil-red O staining and IF were simultaneously executed. Representative images showing LTL (green), THP (red), NCC (red), and AQP2 (red). Black asterisks represent IF-positive tubules. Scale bars, 100 µm.

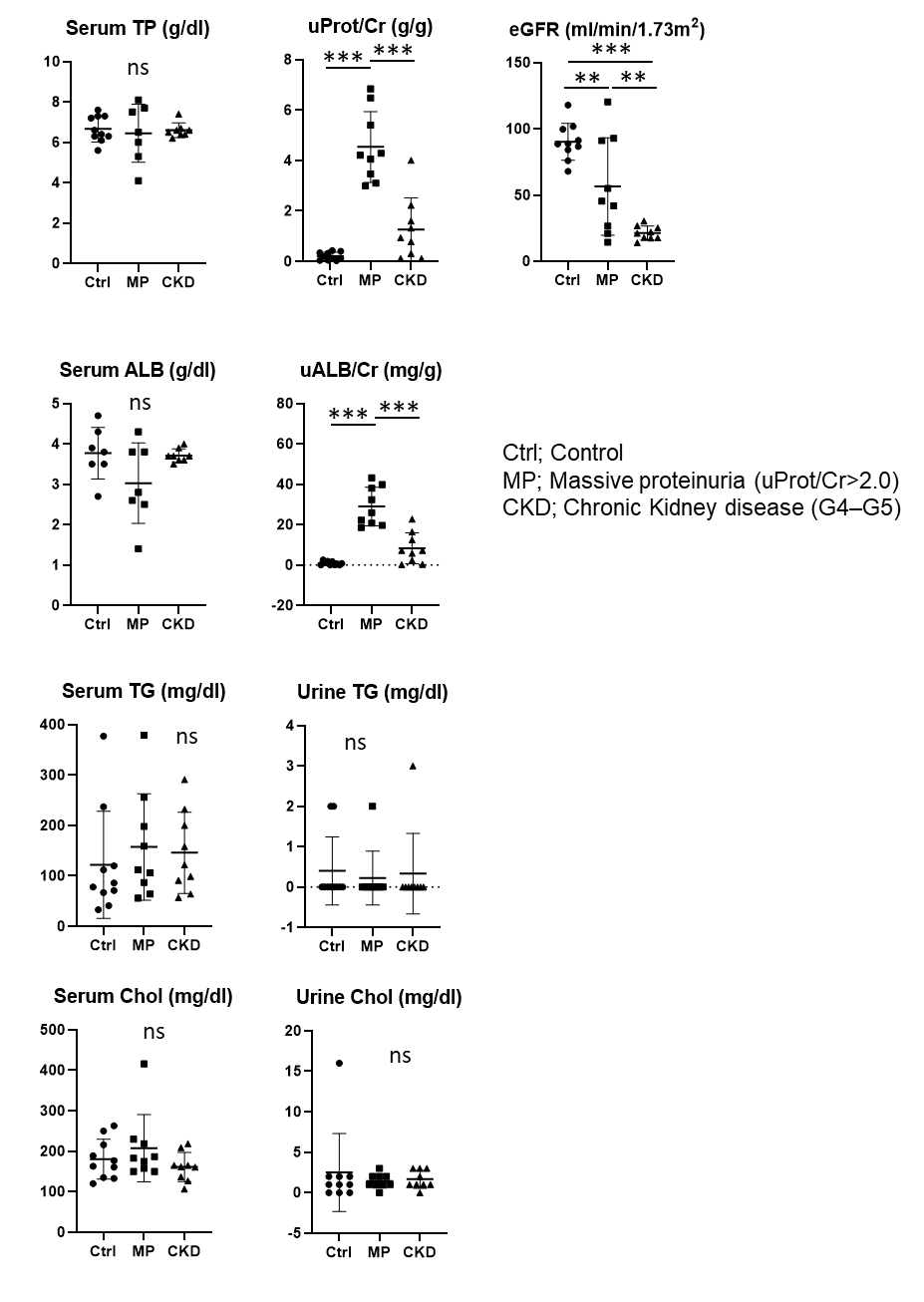

**Figure S6. Biochemical parameters in serum and urine in patients with or without renal diseases.** (n=9–10)
